# Neural coding of temperature with a DNA-based spiking chemical neuron

**DOI:** 10.1101/2022.07.06.498929

**Authors:** N. Lobato-Dauzier, A. Baccouche, G. Gines, T. Lévi, Y. Rondelez, T. Fujii, S. H. Kim, N. Aubert-Kato, A.J. Genot

**Affiliations:** LIMMS, IRL 2820 CNRS-Institute of Industrial Science, The University of Tokyo; Meguro-ku, Tokyo, 153-8505, Japan; Center for Interdisciplinary AI and Data Science, Ochanomizu University; Bunkyo-ku, Tokyo, 112-8610, Japan; Laboratoire Gulliver, UMR 7083 CNRS, ESPCI Paris, PSL Research University; Paris, 75005, France; Laboratoire IMS, UMR 5218 CNRS, University of Bordeaux; Talence, 33400, France; Department of Information Sciences, Ochanomizu University; Bunkyo-ku, Tokyo, 112-8610, Japan

## Abstract

Complex organisms perceive their surroundings with sensory neurons which encode physical stimuli into spikes of electrical activities. The past decade has seen reports of DNA-based chemical neurons that mimic artificial neural networks with chemical reactions. Yet, they lack the physical sensing and temporal coding of sensory biological neurons. Here we report a thermosensory chemical neuron based on DNA and enzymes that spikes with chemical activity when exposed to cold. Surprisingly, this chemical neuron shares deep mathematical similarities with a toy model of a cold nociceptive neuron: they follow a similar bifurcation route between rest and oscillations and avoid artefacts associated with canonical bifurcations (such as irreversibility, damping or untimely spiking). We experimentally demonstrate this robustness by encoding - digitally and analogically - thermal messages into chemical waveforms. This chemical neuron could pave the way for implementing in DNA the third generation of neural network models (spiking networks), and opens the door for associative learning.

**One-Sentence Summary:** A DNA-based chemical network mathematically mimics the sensing of cold by a biological neuron.

## Introduction

Animals represent the world around them with spikes of electrical activities (action potentials) in their nervous system. The spikes are evoked by sensory neurons around the body in response to chemical or physical stimuli (sound, light, pressure, temperature…), and are transported to the central nervous system, where they are decoded, processed and integrated with other sensory signals to elicit a response. A case in point is thermosensation - the sensing of temperature by thermoreceptors and its coding into trains of spikes^1^. Thermosensation can be analog - the firing rates of spikes coding for the ambient temperature^2^. This graded thermal response is essential for organisms which cannot regulate their body temperature and resort to thermal acclimation or thermotaxis to adapt and navigate the thermal constrains set by their environment ^3^. Thermosensation can also be digital like in thermal nociceptors, where temperature-gated ion channels such as TRPM8 receptors generate trains of spikes only when they are exposed to dangerous levels of hot or cold^1^, and remain at rest otherwise (Fig 1A). This neural coding of sensation is sparse in space and time, allowing a fast and accurate response to be computed from a few transient signals^4^.

**Figure 1:**
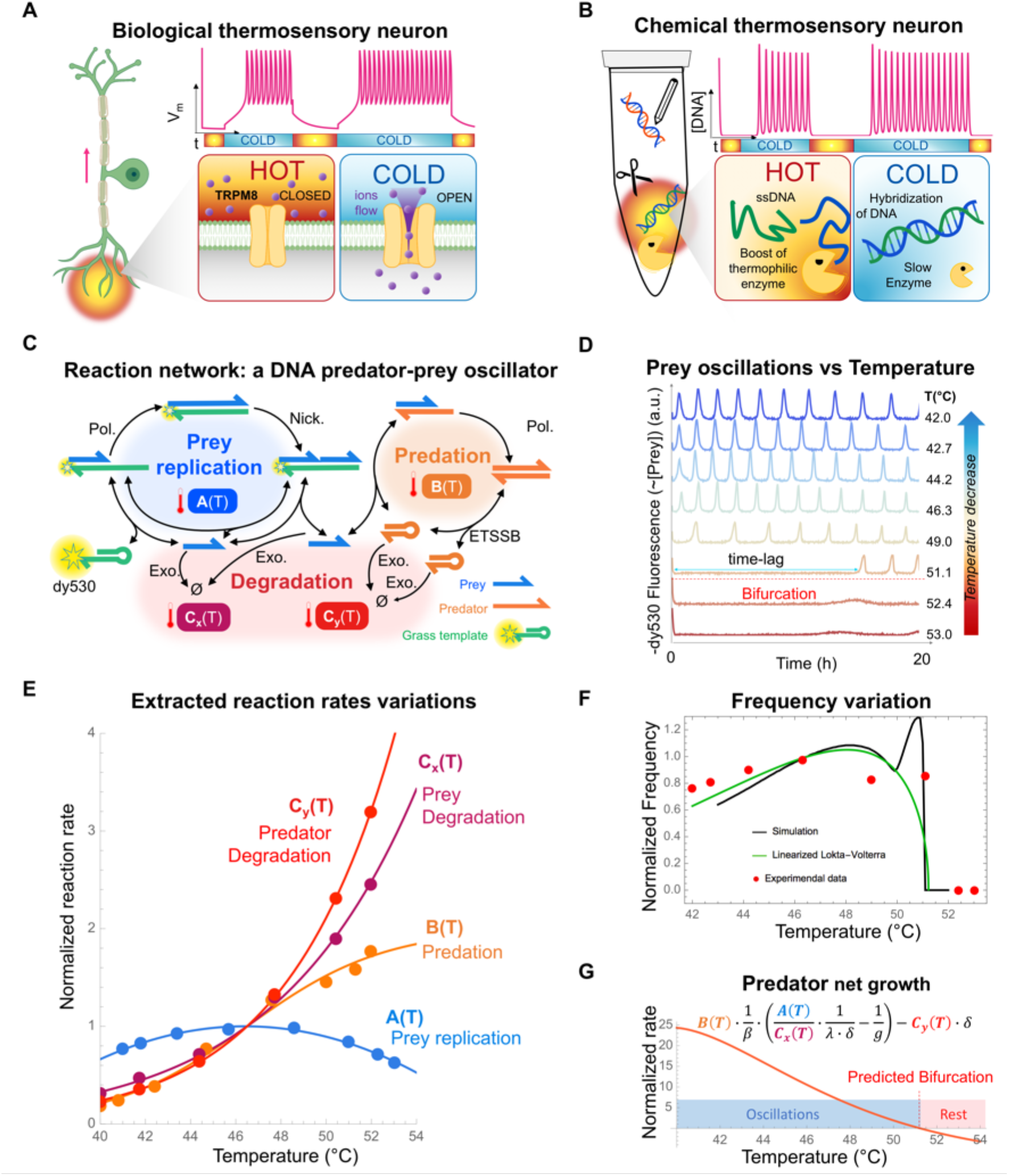
Cold sensation by a chemical neuron. (**A**) A cold sensory biological neuron switches between a rest state at high temperature to a spiking state at low temperature. The bifurcation is controlled at the molecular level by temperature-sensitive ion channels. (**B**) Our cold sensory chemical neuron switches between a rest state at high temperature to a spiking state at low temperature. The bifurcation is controlled at the molecular level by temperature-sensitive enzymes and the melting of DNA duplexes. (**C**) The chemical neuron is based on a DNA-based predator-prey oscillator, which comprises three reaction modules (prey replication, predation and degradation) that are actuated by three DNA-processing enzymes. (**D)** Empirical dependence of the oscillator on temperature. The fluorescent traces are measured in bulk and mainly reflect the level of prey. (**E**) Individual measurements of the reaction rates of the three modules with temperature (SI 2.2). For sake of comparison, rates are normalized by their value at 46.5 °C. (**F**) Predicted and measured frequency of spiking against temperature. The black plain line shows the prediction of a nonlinear enzymatic model, and the green plain line shows the prediction its linearized version (Lotka-Voltera model) (SI 2.3). The red dots show the experimental measurements. (**G**) Prediction of the net rate of predator growth in the rest state (low predator) (SI 2.4). This rate includes the effect of predation, prey growth and predator decay. When the net rate is negative, the rest state is stable. When the rate becomes positive, the system bifurcates the rest state disappears and oscillations emerge.

In electronics, the parsimony and efficiency of neural coding has motivated the development of Spiking Neural Networks, which are electronic oscillators connected by a network of excitatory and inhibitory synapses^5,6^. Departing from the heavily supervised, energy and data hungry paradigm of deep neural networks, spiking neural networks can in principle learn and operate much more frugally^7,8^: with few neurons and with little supervision, data or power – similarly to biological neurons^4,9^.

In chemistry, previous works has hinted at the feasibility of programming nonlinear chemical reactions to emulate neurons^10–12^ Emboldened by the success of DNA as a building^13^ and computing^14–16^ material, various groups have reported DNA-based neurons capable of performing linear^17,18^ and nonlinear classification^19,20^ on nucleic acids. However, these DNA neurons still pale in comparison with their biological counterparts. Firstly, they only process chemical stimuli (concentrations of nucleic acids), and are unable to integrate physical stimuli such as temperature. Secondly, these DNA neurons are akin to a toggle switch and have no sense of time. They are incapable of processing information in the time domain (e.g. deciding if two inputs are present sequentially or simultaneously), and lack the temporal finesse of biological neurons, which encode their stimulation history into a spike waveform^21^. With this temporal coding, the brain can decide if two stimuli are sequential or simultaneous (by comparing the arrival time of spike trains^22^), and learn from this temporal correlation to wire together neurons that fire together. This is the basis of associative learning, which allows the brain to store a sparse representation of its environment in its neural topology^23^. Chemical systems that mimic this temporal coding would open the door for associative learning and enables a novel range of molecular systems that learn from their environment. Encoding information with temperature also has practical benefits: it is easy to implement with a thermal cycler^24^, and the modulation of temperature to probe and program molecular systems has been studied for more than a decade^25–29^.

Here we report a thermosensory chemical neuron which encodes thermal signals in the temporal domain. The chemical neuron produces spikes of chemical activity when exposed to low temperature (Fig. 1B) - similarly to a cold nociceptive neuron^30,31^. At high temperature, the chemical neuron is at rest and does not spike. As temperature is lowered and crosses a threshold, the neuron bifurcates to an oscillatory regime, turning into a chemical metronome that steadily generates pulses of a DNA species - the firing rate coding for the ambient temperature.

Comparing our chemical neuron with a toy-model of a thermosensory biological neuron, we find that they bifurcate along a similar route. This route reversibly and cleanly switches the neurons between rest and oscillation, avoiding artefacts of simpler bifurcations (such as irreversibility, damping or erroneous spiking). Building on this robustness, we operate the chemical neuron in bulk as a digital communication channel –demonstrating high fidelity and resilience to thermal noise. Lastly, we miniaturize the chemical neurons and operate tens of thousands of them in an analog mode, precisely coding the ambient temperature into the firing rates of the neurons. Such chemical neuron opens an avenue to execute recurrent molecular computations and connect chemical neurons into a spiking chemical network, which could process information in a manner similar to the nervous systems of complex organisms.

### The spike generator

Spiking neurons can be conceptualized as oscillators that produce rhythmic pulses of activity^32^. We selected a predator-prey chemical oscillator^33^ to generate chemical spikes. Although this DNA oscillator is simple, it is one of the most robust reported^34–40^ and oscillates for days on end^41^. It can be wired to other DNA circuits to cadence their activities, for instance to rhythmically aggregate and disaggregate colloids ^42^. This enzymatic oscillator comprises two DNA strands, a prey and a predator, which live and die according to rules set in their sequences and enforced by 3 enzymes (polymerase, nickase, exonuclease). Briefly, a prey replicates by binding to a grass template, an event which recruits a polymerase and a nickase and produces a new prey as an output (Fig. 1C). This self-replication produces an exponential growth in the population of prey (prey growth). At the same time, the predators replicate by “eating” the preys, converting them into predators through an enzymatic reaction that produces one new predator for each prey consumed by a predator. This predation triggers an exponential growth of the population of predators at the expense of preys, whose population then decays exponentially – eventually choking the growth of predators themselves. Once the exonuclease has cleared the population of predators, the remaining preys resume their exponential growth, and the cycle of growth and predation restarts - resulting in rhythmic oscillations with a typical period of ∼1 h.

We first took a phenomenological approach and investigated how this oscillator depended on temperature. We found that it behaves similarly to a cold sensory neuron - being inactive over a temperature threshold and suddenly becoming active below (Fig. 1D). At 52.4 °C or above, oscillations are silenced and the system is locked in a stable steady state with a low level of preys. But as temperature is lowered near 51°C, the system suddenly bifurcates and enters an oscillatory regime. At 51.1°C, oscillations emerge after a lag of dozens of hours. At 49°C, oscillations quickly kick in and the system adopts a rhythmic pattern. The firing rate varies non-monotonically with temperature: it increases from 49°C to 46.3 °C, at which point it decreases (Fig 1F). By contrast, the amplitude of oscillations remains comparatively stable with temperature.

Although the oscillator bifurcates between rest and oscillations in a manner similar to a cold sensory neuron, this phenomenological approach fails to precisely explain why and how it does so. The net effect of temperature on oscillations is difficult to predict from the chemical network of the oscillator, as temperature affects the thermodynamics and kinetics of all chemical reactions. Heating promotes the melting of DNA duplexes, accelerating the separation of predators (involved in predator growth) and the melting of prey from their template (involved in prey growth). But heating also destabilizes the binding of predators to prey, and the binding of preys to their templates. Increasing temperatures also affects the kinetic balance of enzymes, and thermophilic enzymes like the exonuclease are expected to become more active with increasing temperature.

We thus took a reductionist approach to understand the kinetic basis of why and how the chemical neuron bifurcates with temperature. We started from a validated mathematical model of the oscillator^33^, which is two dimensional (tracking the dynamics of prey and predators) and nonlinear (accounting for the saturation of enzymatic kinetics through Michaelis-Menten equation), but which lacks temperature dependencies (SI 2.1). We isolated each of the three reaction modules of the model (prey growth, predation, degradation of prey and predators) and quantified their temperature dependence (Fig 1E & SI 2.2). Prey replication varies weakly (and non-monotonically) with temperature. By contrast, predation accelerates more than 5-fold from 40 °C to 54 °C – suggesting that the separation of predators - promoted by heating - is the rate-determining step in predation. Lastly, the degradation of prey and predators speeds up more than 10-fold over this temperature range - which is consistent with our exonuclease being thermophilic^43^. We validated this modelling with a *a priori* prediction of the bifurcation temperature of Figure 1D. We linearized the enzymatic saturations - reducing the model to the celebrated Lotka-Volterra equation, and used classical linear stability analysis to express in a closed-form the frequency of oscillations and their temperature of bifurcation. Satisfyingly, this simplified model predicts the actual bifurcation temperature within ∼1 °C (Fig 1F & SI 2.3).

Coming back to the enzymatic model, we sharpened our understanding of the molecular basis of temperature-induced bifurcations. We postulated that oscillations emerge once the steady state at high-temperature loses its stability (SI 2.4). In this steady state, the level of predators is low, and the level of preys is mainly set by the balance between their replication and degradation by the polymerase – which essentially reduces the dynamics of predators to a one dimensional linear different equation. Knowing the steady-state level of prey, we computed the net growth rate of predators as the outcome of two linear processes: replication by predation of the prey and degradation by the exonuclease. At high temperatures, the exonuclease is highly active and there are not enough preys to feed the predators and counter their degradation by the exonuclease. The net growth of predators is negative and the steady state remains stable. But as temperature decreases, the exonuclease slows down, which has compounding effects on the growth of predators. First, it raises the steady-state level of preys, which promotes the replication of predators by predation, and it also attenuates the degradation of predators. Overall, the model predicts that replication and degradation cancel out at ∼51°C (Fig. 1G) – a theoretical prediction that agrees within less than 1°C with the temperature measured experimentally for the bifurcation (Fig. 1D).

In addition, we confronted the full model (2D and nonlinear) to an experimental bifurcation diagram-varying both temperature and exonuclease. The model satisfyingly predicts the onset on experimental bifurcations when those two parameters are varied. (Fig. S5)

### Bifurcation routes

Comforted by the predictive power of model, we visualized the bifurcation route of the neurons from rest to oscillation (Fig. 2A). We consider the prey nullcline (green) and predator nullcline (blue). At high temperature, trajectories are attracted to the stable steady state with few predators (black point near the *x* axis) and the neuron rests. As temperature is lowered, the steady level of prey increases, dragging the tip of the prey nullcline to the right of the plot. When the tip intersects the predator nullcline, a first saddle-node bifurcation occurs - creating an unstable state and a stable state, which quickly transitions to a stable spiral state - generating damped oscillations in its vicinity. Yet a generic trajectory (pink curve) cannot reach this spiral state because it is attracted by the stable state with low predators, and this saddle-node bifurcation is not apparent from a generic time trace. But as temperature decreases, the unstable and stable states keep sliding along the nullclines and eventually annihilate each other. This is the situation mentioned above where the net growth rate of predators enters the positive zone (i.e. the region on the right of the predator nullcline). This second saddle-node bifurcation clears the way for a generic trajectory to reach the spiral. It manifests in the time trace as damped oscillations that start after a long time-lag, which is due to the remnant of the stable state (i.e., the growth rate of preys and predators near this point is positive, but still close to 0 by continuity). Lastly, as temperature decreases again, a Hopf bifurcation destabilizes the spiral state and gives birth to a limit cycle. This manifests in the time trace as sustained oscillations that set in quickly from generic trajectories. The 3 bifurcations occur in a narrow temperature range (∼0.1°C) (Fig. 2A & SI 2.6). While the ordering of the last two bifurcations (the second saddle-node and Hopf bifurcations) is sensitive to parameters, the general bifurcation path (destruction of a resting state and creation of a limit cycle) remains the same.

**Figure 2.**
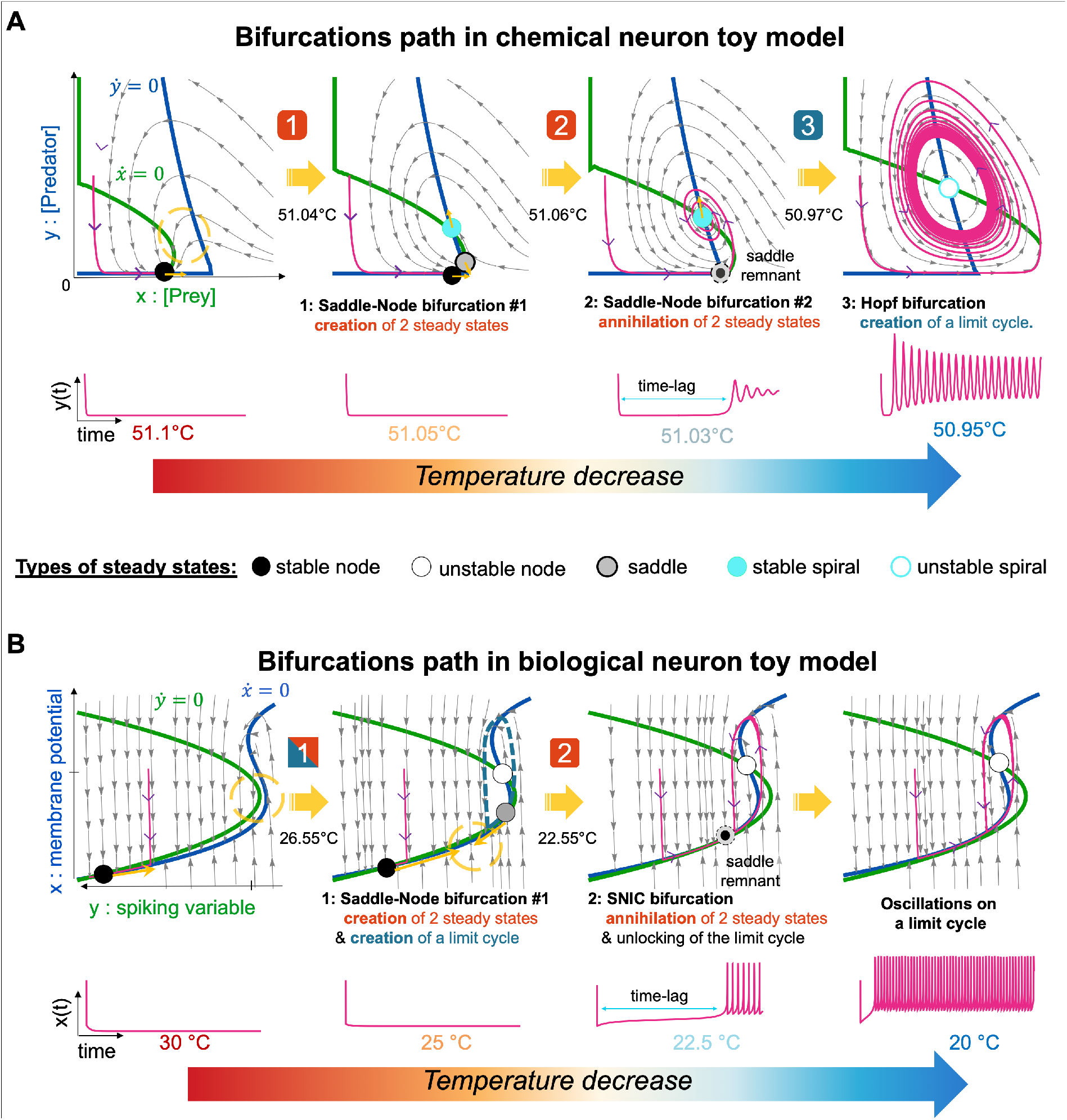
Bifurcation routes of our chemical neuron and a toy model of a thermosensory biological neuron. (**A**) 2D phase portraits of the chemical neuron for decreasing temperatures (see Movie S1). Thick green and blue lines are the nullclines for preys and predators respectively. The thin pink line shows a generic trajectory. Steady states – located at the intersection of nullclines – are shown as disks, color-coded by their nature and stability (see SI 2.6) (**B**) 2D phase portraits of a toy model of biological thermosensory neuron (see Movie S2), which has 2 variables (membrane potential and spiking variable), and is parametrized by temperature through its excitation current (see SI 3.1).The blue curve is the nullcline of the membrane potential, and the green curve is the nullcline of the spiking variable.

We then compared our chemical neuron with a toy model of a cold sensory neuron. While the mechanical basis of temperature sensing in neurons has been known for more than a decade, only a few mathematical models have attempted to explain how this temperature-dependence governs the bifurcations between rest and spikes.^44–46^ These models are typically based on a Hodgkin-Huxley-type model^47^ (with a thermal dependence of the current to model the influence of temperature on thermoreceptors). Yet the Hodgkin-Huxley model is four dimensional, which makes the visualization and analysis of bifurcations delicate. Another model, the FitzHugh-Nagumo model, simplifies the dynamic of spiking by reducing the number of variables from four to two, which makes it analytically tractable and easier to visualize ^48–51^. Yet, this model is limited to describing a certain class of neurons (those with class 2 excitability) due to the linear shape of one its nullcline (the nullcline for the recovery variable, also called spiking variable).

For the sake of generality, visualization and comparison with our chemical neuron, we selected a generalization of the FitzHugh-Nagamo model with a quadratic nonlinearity in the spiking variable^52^ (which makes it mathematically equivalent to the 2D Hindmarsh-rose model^49^). Being 2D, this model retains the ease of visualization and analysis of the FitzHugh-Nagamo model, while covering more classes of neuronal excitability thanks to its quadratic term. The model features only two variables (the membrane potential *x* and the spiking variable *y*), and four parameters (of which only three are freely tunable, because changing the parameter *a* has the same effect as changing the current). We set two parameters (*c* and *d*) to fall in a regime where bifurcations have been analytically studied in details (corresponding to a Class 2 excitability and Class 1 spiking^52 53^). We fix the remaining free parameter *b* to 0.6 to analyze and plot the bifurcations, and compare them with our chemical neuron (but the conclusions remain broadly similar with other values of *b*). Lastly, to capture the effect of temperature, we added a sigmoidal dependence of the excitation current *I* on the temperature *T*, which qualitatively agrees with the shape of measurements of *I(T)* in thermosensitive ion channels such as TRPM8^54^ (SI 3.1)

Since this model is a toy model, it does not pretend to describe how any chemosensory neuron will bifurcate with temperature. But it describes a plausible route for bifurcations, a route that is generic, analytically tractable and easy to visualize– allowing a qualitative and visual comparison with our chemical neuron. In this route, at high temperature the biological neuron is at rest, attracted by a stable steady state (Fig. 2B). As temperature decreases, the tip of the *y* nullcline intersects the *x* nullcline – a saddle-node bifurcation that creates two steady states: one stable and one unstable. It also creates a limit cycle which is blocked by the saddle that lies on it. Lowering the temperature again, a SNIC (Saddle Node on an Invariant Circle) bifurcation occurs that annihilates the rest state and saddle. This clears the way for the biological neuron to reach its limit cycle from a generic point in the phase space.

The biological and chemical neurons thus follow a similar route from rest to oscillation (or from oscillation to rest), by destroying (creating) their rest state and creating (destroying) the limit cycle that supports their oscillations. This sequential bifurcation route offers several advantages for spike-encoded sensing compared to more traditional bifurcations (Fig. 3 & SI 3.2). Among the canonical bifurcations between rest and oscillations (Saddle-Node, SNIC, homoclinic, supercritical and subcritical Hopf, Fold Limit cycle), only the SNIC and supercritical Hopf bifurcation are reversible – in the sense that they do not suffer from hysteresis when the control parameter is repeatedly swept across the bifurcation^55^

**Figure 3.**
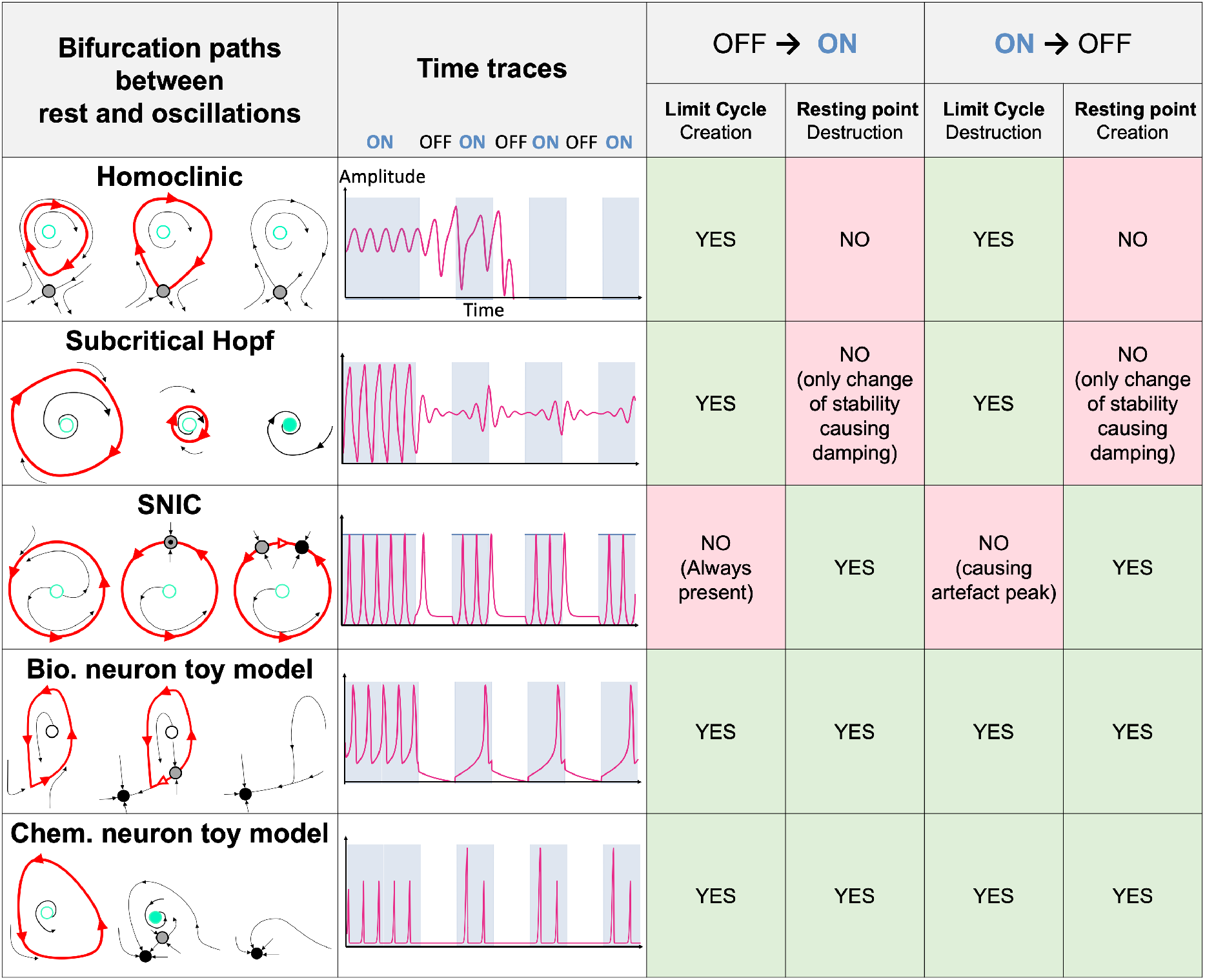
Response of canonical dynamical systems, and chemical and biological neurons to periodic stimulations. These dynamical systems are subjected to periodical switching of their bifurcation parameter (SI 4), causing bifurcations between a rest state (OFF) and a limit cycle (ON). Some systems bifurcate by creating and destroying their rest state (e.g., SNIC), others by creating and destroying their limit cycle (e.g. Hopf). But only the biological or chemical neurons create and destroy both their limit cycles and their resting state. This allows them to switch between spiking and rest state without artefacts and reversibly (contrary to the canonical bifurcations).

Similar to SNIC and Hopf bifurcations, our chemical and biological neurons are reversible and do not exhibit hysteresis. This is because their rest state and limit cycle do not coexist – except within a narrow temperature range. Outside this temperature range, the state of the neuron is unambiguous and only determined by temperature – not by the history of the neuron or its stimulation. This prevents the memory effect or hysteresis which occurs with say homoclinic bifurcation (SI 4.1).

Like in a Hopf bifurcation, the chemical and biological neurons create and destroy their limit cycles. But unlike a Hopf bifurcation, they enter or leave the oscillatory regime in a clean and undamped manner because they create and destroy their steady states – removing the spiral state that pollutes the transitions between rest and oscillations (SI 4.2). Since steady states can only typically be created or destroyed in pair, this route needs an auxiliary saddle steady state, which is used to destroy the rest state (when temperature is lowered and the system is switched from OFF to ON), or to destroy the steady state in which the limit cycle is shrunk (when temperature is raised and the system is switched from ON to OFF).

In other words, the bifurcation route of the chemical and biological neurons is hybrid and combines the destruction (creation) of a stable node with the creation (destruction) of a limit cycle. With this combination, the neurons robustly sense temperature – reversibly and cleanly switching between ON and OFF states. This is confirmed by numerical simulations of toy models of canonical bifurcations (see SI 4): Homoclinic, Hopf and SNIC bifurcation are subject to artefacts when their control parameter is repeatedly turned ON and OFF, while the chemical and biological neurons do not suffer from such artefacts (Fig. 3). We experimentally verified this robustness for the chemical neuron, and we additionally confirmed that it resisted severe thermal noise.

### Passing of thermal messages

Equipped with this theoretical grounding, we operated our neuron as a coding channel and encoded digital profiles of temperature (thermal messages) in which the neuron is switched between ON and OFF states. Each thermal message was divided in ten intervals of time, each interval carrying one bit of information (Fig 4A). During each interval, the temperature is set to either hot if the bit is 0 (which causes the neuron to rest, being OFF), or cold if the bit is 1 (which causes the neuron to spike, being ON).

**Figure 4.**
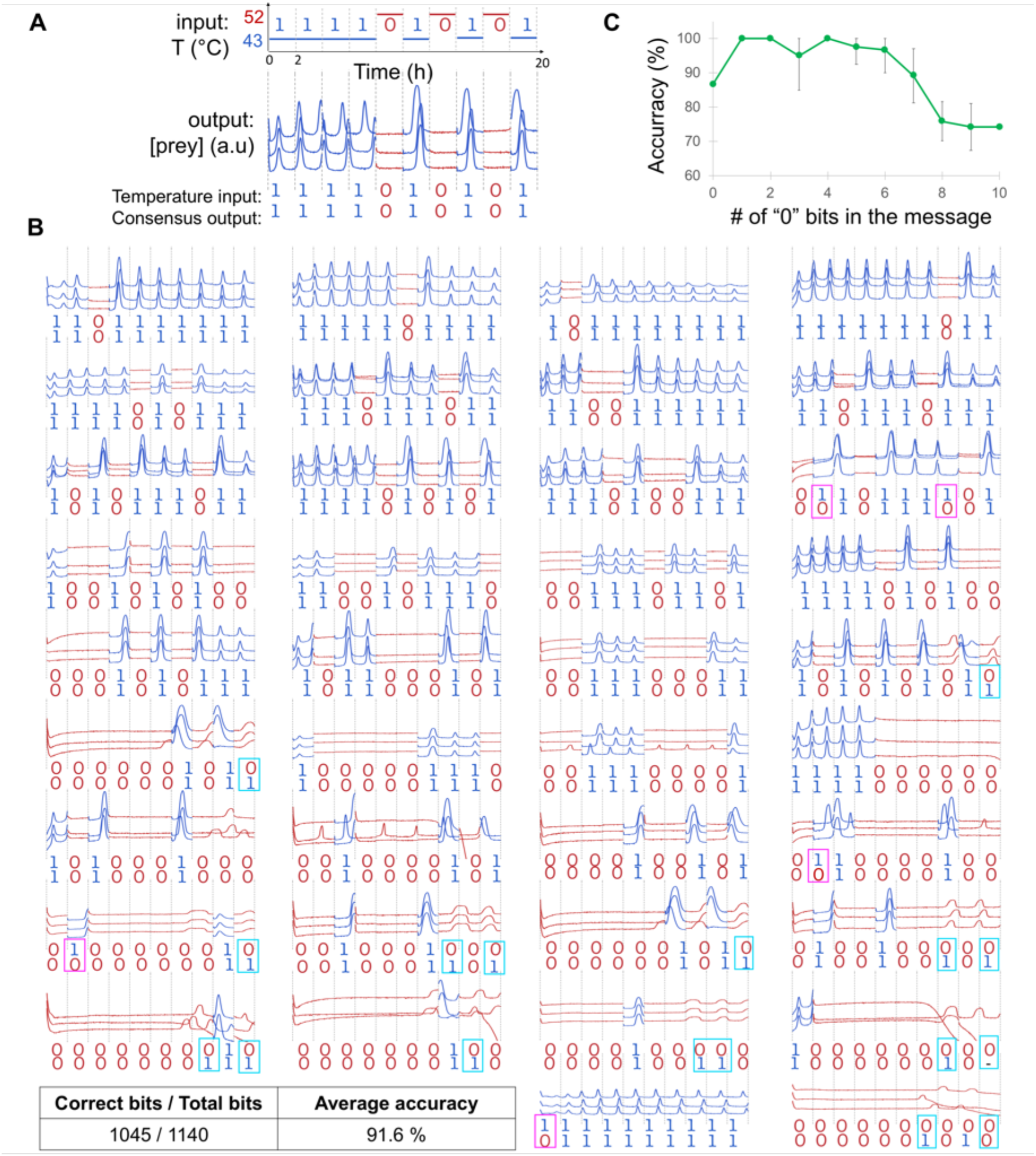
Modulation of 10-bits spike trains with thermal messages. (**A**) The predator-prey system is subjected to various digital profiles of temperature. For each interval, the temperature is set either to hot (52°C) if the bit is 0, or cold (43°C) if the bit is 1. Experiments are performed in triplicate and the fluorescence of the system tracked. The consensus output is the consensus sequence obtained by majority voting on the three replicates. Erroneously transmitted bits are highlighted in light blue. (**B**) Time traces of 38 random messages. Messages are grouped according to their number of 0 bits. Magenta and cyan frames denote respectively an error on a 1 bit or a 0 bit. (**C**) Accuracy of transmission against the number of 0 bits in the message. The accuracy for each group is measured (defined as the number of correct bits transmitted, averaged over all bits of all the triplicates of all the message of a given group).

We first optimized the hot and cold temperatures (SI 5). The hottest temperature achievable is dictated by the emergence of parasites, which are DNA species that have evolved to resist degradation by the exonuclease, and which poison the system by monopolizing enzymatic resources ^56^. While the precise mechanism for the emergence of parasites is unclear, it is activated by temperature and is thought to involve breathing and partial melting of DNA strands.

We thus set the hot temperature (52°C) for the OFF state, only slightly above the bifurcation temperature (∼51°C) to mitigate the emergence of parasites. On the other side, the cold temperature (ON) was dictated by the timescale of our experiment. We found that below 43°C, the priming of spikes was too slow to be practical and we chose this temperature as the cold temperature. Noticeably the hot and cold temperature are not symmetric with respect to the bifurcation.

After optimization, we passed thermal messages of 10 bits. Experimental results confirm theoretical insights from the model: the neuron does not damp when temperature is switched (unlike Hopf bifurcations), and it does not spike extraneously when turned OFF (unlike SNIC bifurcations). The state of the neuron mostly depended on the inputted temperature, not its history (Fig. 4B). More than 90% of the bits were correctly coded (Fig. 4C) and most errors were false positives (e.g., spikes at high temperature, light blue boxes in Fig.4) that occurred after long stretches of 0s during which the neuron remained at high temperature for a long time. This is consistent with the view that parasitic species emerge at high temperature and interfere with the normal functioning of the neuron. Working at the microscale could mitigate the emergence of parasites, since it is a stochastic event that scales with the volume of the system. Compartmentalization in micrometric droplets has been shows to suppress parasitic species in other species,^57,58^, and it could drastically extend the length of messages that can be passed in our system.

In a separate experiment, we further challenged the neuron by shrinking the thermal amplitude (the difference between the hot and cold temperature) and adding thermal noise (Fig. 5). Rather than keeping the cold or hot temperature constant, we swept it around the mean hot or cold temperature, adding noise to the input temperature. For a cold temperature of 46 (+/-2) °C and a hot temperature of 49 (+/-2) °C (giving a signal to noise ratio of 3°C/2°C=1.5), the neuron was still able to encode a message of 5 bits. Such a thermal window is sufficiently narrow so as not to perturb significantly downstream processes, and would allow the integration of the neurons with DNA-based logic circuits or neural networks^19^.

**Figure 5.**
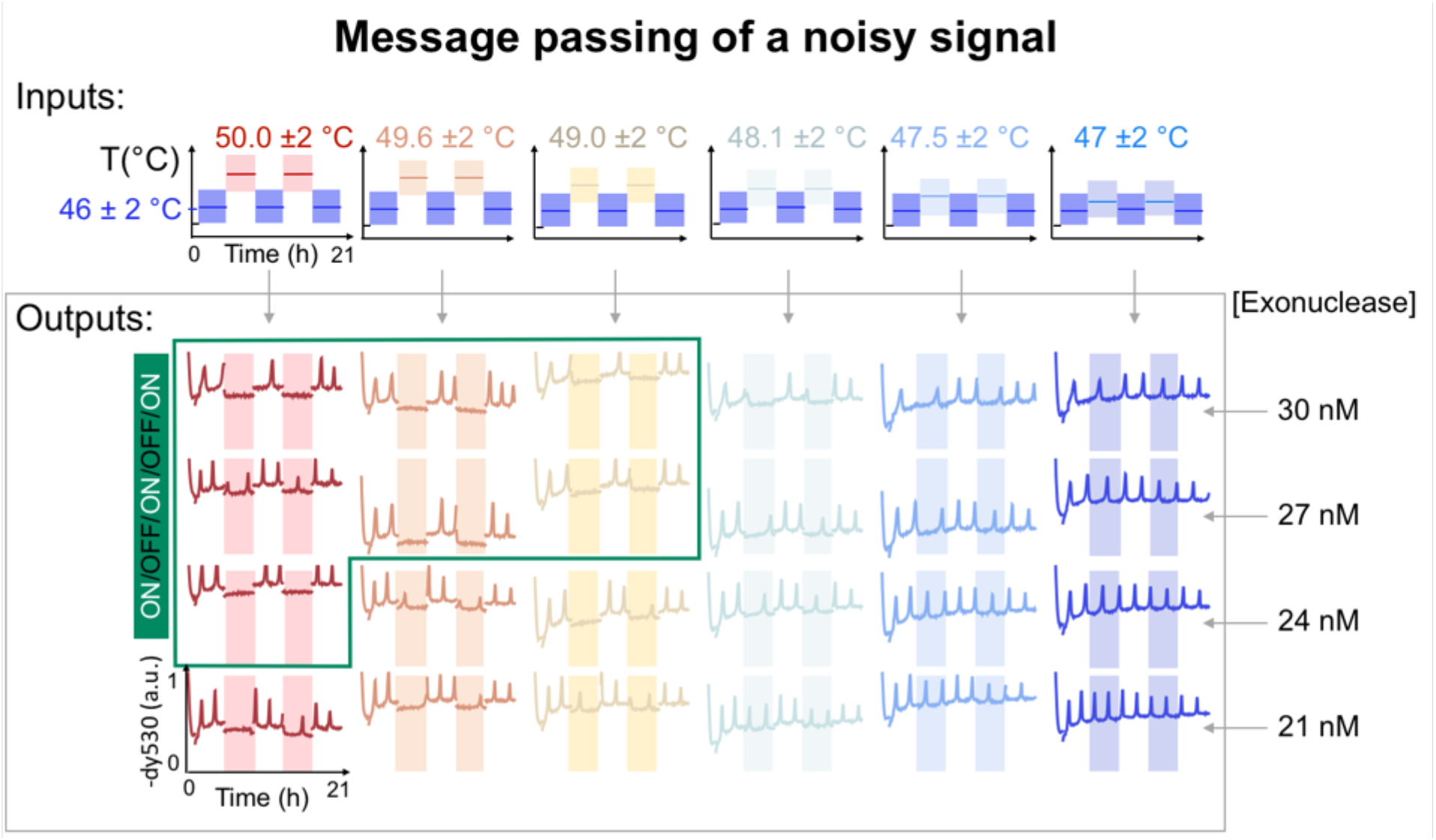
Response of the chemical neuron to noisy temperature stimuli. During each interval, the temperature oscillates 2°C above and below the nominal temperature with a 3 min period. The zone corresponding to successful passing of the message is highlighted in green with the exception of the 50°C/27nM message which contains some errors.

### Analog encoding of temperature

After operating the chemical neuron in a digital mode, we asked if it could be operated analogically, i.e. encoding the ambient temperature in the spiking frequency? This analog mode of operation is used by some organisms to track changes in temperature, enabling for instance thermotaxis.

To that end, we miniaturized the chemical neuron and mapped *en masse* its dependence on temperature and exonuclease. Briefly, we prepared ∼30,000 micrometric droplets containing the chemical mix and varying concentrations of exonuclease, which we incubated in a silicon chamber^59^ (a material with excellent thermal properties) placed in a thermal gradient (the temperature range being selected to fall in the oscillatory regime) (Fig. 6A). This microfluidic mapping reveals a temperature range where the firing rate of spikes varies linearly with temperature (Fig. 6B-C), and where the amplitude of spikes remains roughly constant (Fig. 6E-F). This range is ideal to operate the chemical neuron as a temperature sensor, as it mimics the linear encoding of temperature seen with some thermosensory neurons^2^. Interestingly, temperature and exonuclease mostly compensate each other: the same firing rate can be obtained by compensating an increase in temperature by a decrease in the concentration of exonuclease, which facilitates the adjustment of parameters.

**Figure 6.**
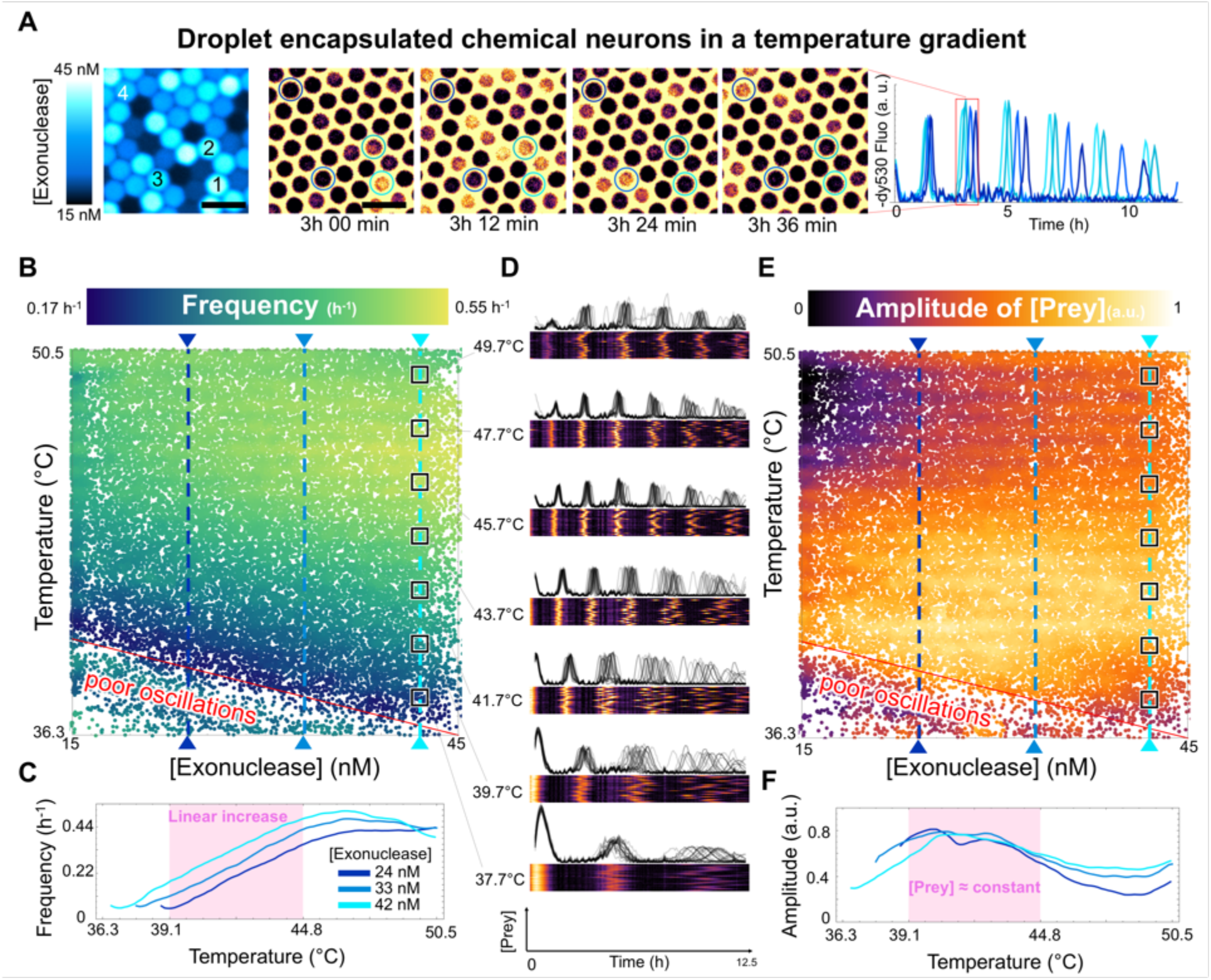
[Exonuclease]-Temperature phase diagrams of the chemical neurons. (**A**) Time-lapse of encapsulated chemical neurons in a temperature gradient (movie S3). Varying the concentration of exonuclease - given by a fluorescent barcoding (left and Image S1) – changes the frequency of the oscillations. scale bar: 100 μm (**B**) Experimental frequency phase diagram. The zone corresponding to low temperature and low concentration of exonuclease is populated of neurons that oscillate poorly (< 3 oscillations). For [Exonuclease] >30 nM the variation of frequency with increasing temperature is non monotonous and a maximum can be observed around 47°C. (**C**) In the 39°C - 45°C range the frequency increases linearly with temperature and its level can be offset by varying the exonuclease. (**D**) Peristimulus Time Histograms at different temperatures. In the 46°C - 48°C, droplets are very synchronized compared to droplets at lower temperatures. (**E**) Experimental amplitude phase diagram. (**F**) Evolution of prey amplitude with temperature for 3 concentrations of exonuclease.

We plotted Peristimulus Time Histograms (PSTH) to compare the oscillation of droplets with the same content and incubated at the same temperature (which is inspired by PSTH diagrams for biological neurons exposed to the same stimuli). Like their bulk counterpart, the chemical neurons in droplets oscillate rhythmically (Fig. 6D). But micrometric compartmentalization reveals features that are invisible in bulk, namely stochastic effects. At high temperature, the droplets oscillate synchronously, but at lower temperature, they progressively desynchronize (which is visible from the spreading of the spikes). We posit that this desynchronization likely emerges from the low copy number of species in droplets. When the concentrations of prey reach the sub picomolar range, their copy number goes below ∼100 copies, and their chemistry cannot be correctly modeled with deterministic equations. Rare events may be amplified by the nonlinear chemistry, giving rise to large deviations from the deterministic equations. For instance, a template (which is normally in a close hairpin conformation) may transiently open up, allowing a prey to bind and replicate. Or two preys that bind fleetingly may be extended by the polymerase, giving rise to two predators that replicate exponentially. Enzyme fatigue (i.e. the partial unfolding and loss of activity of enzymes) accelerates at higher temperature. This may explain how droplets in the 45-48 °C range remain synchronized.

Overall, this shows that in principle the thermal history of the neuron could be recorded into the temporal profile of its chemical spikes and reconstructed by playing back the time course and measuring how the firing rate of spikes change with time.

### Discussion

We have built a chemical neuron that spikes with chemical activity when exposed to cold – similarly to a cold sensory neuron. At a fundamental level, this chemical coding is temporally sparse: the signal is localized in time because the neuron only activates when stimulated. This reduces power consumptions and opportunities for chemical “cacophony” (*i*.*e*. cross-talks between unrelated chemical systems). Sparse encoding also allows spikes to be compared and sorted according to their timing and frequency, which forms the basis of information processing in the brain ^4^.

The dynamics of a single chemical neuron could already be rich enough to support advanced computations like reservoir computing ^24,60^. In this paradigm, a chemical neuron would be programmed to accept a time-varying signal as an input, and return for instance the time-lagged or time-averaged version of this signal as an output. More generally, the neuron could be programmed to fire when it recognizes a specific temporal pattern (like the alternance of high and low signals at a given frequency).

Yet the full benefits of temporal coding will only be unlocked once neurons are connected into a network. For instance, warmth could be represented by integrating signals emanating from cold and hot sensory neurons (similarly to the biological sensation of warmth ^30^). Here we encapsulated millions of neurons in vesicles with microfluidics, but they remain to be wired with chemical synapses that traffic DNA between vesicles. Water-in-oil droplets are impermeable to DNA, but semi-permeable vesicles, or liposomes equipped with pores ^61,62^) could form the basis of such chemical synapses.

Once the chemical neurons are connected together, their weights (i.e. the rate of chemical exchange between two neurons) could be learned by chemically reinforcing links between neurons that spike synchronously when exposed to the same stimuli. This Hebbian learning rule (“neuron that fire together wire together”) is already used to learn weights in electronic spiking networks ^23^.

Chemical networks of thermosensory neurons could be integrated with locomotion to perform thermotaxis, by detecting change in the frequencies (i.e., the ascent of descent of a thermal gradient) and adjusting locomotion accordingly. More generally, networks of spiking chemical neurons could perform phototaxis, chemotaxis or tasks which require the integration of sensing, computation and locomotion at the molecular scale.

## Supporting information

SUPMAT-Neural coding of temperature with a DNA-based spiking chemical neuron

READ ME - Videos & Image Legends

Movie S1-ChemNeuron_Bifurcation_Path

Movie S2-BioNeuron_Bifurcation_Path.mp4

Movie S3-Encapsulated Chemical Neurons in a Temperature gradient

Image S1-Barcoding of Exonuclease concentration in Encapsulated Chemical neurons

## Acknowledgments

We thank Kazuyuki Aihara, Takashi Kohno, Ibuki Kawamata, Yusuke Sato, André Estevez-Torres and Jean-Christophe Galas for discussion on the manuscript. We thank FEMTO-ST (CNRS, Besancon) for the fabrication of Sillicon microchambers. Neurons and ionic channels in Figure 1 were adapted from BioRender.com

## Funding

JSPS KAKENHI Grant Numbers JP19KK0261 (NAK), JP20K12061 (NAK, AG)

CNRS MITI interdisciplinary program on Biomimetism (GG, TL, AG).

## Author contributions

Conceptualization: NLD, AB, GG, TL, YR, NAK, AG

Methodology: NLD, AG

Investigation: NLD

Visualization: NLD

Funding acquisition: GG, TL, NAK, AG

Supervision: TF, SHK, NAK, AG

Writing – original draft: NLD, NAK, AG

## Supplementary Materials

Materials and Methods

Supplementary Text

Figs. S1 to S17

Tables S1

References (*1*–*18*)

Movies S1 to S3

Image S1

## Notes

### Competing Interest Statement

The authors have declared no competing interest.

### Summary of Updates

Figure 2 and corresponding text revised Supplemental files updated Addition of movies S1-S3 and Image S1

