## Supplementary material for "Neural coding of temperature with a DNA-based spiking chemical neuron": SUPMAT-Neural coding of temperature with a DNA-based spiking chemical neuron

### 1. Experimental methods

#### 1.1. Oligonucleotides

The prey and predator strand were purchased from Integrated DNA technology (IDT) and the template G1 was purchased from Biomers (Germany). All strands were suspended in 1X Tris-EDTA buffer (Sigma-Aldrich) containing 1 M Tris-HCl and 0.1 M EDTA at 100 µM. The template G1 was protected with phosphorothioate backbones on its 5' end to prevent degradation from the exonuclease. The sequences are listed in Table 1. They correspond to the predator-prey in [1]. Template G1 was diluted to a stock concentration of 5 µM, and used in a range varying from 45 to 90 nM (0.9 % to 1.8 % of the working solution). In most experiments, prey and predator strands were diluted to 1 µM and were used in a range varying from 1 to 10 nM each (0.1 % to 1% of the working solution).

| **Name** | **Sequence** | **5' modification** | **3' modification** |
|---|---|---|---|
| Template G1 | C*G*G*CCGAATG-CGGCCGAATG | | dy530 |
| Prey N1 | CATTCGGCCG | | |
| Predator P1 | CATTCG GCCGAATG | | |
| **Note** | * phosphorothioate backbone | | |

**Table S1 - PEN Predator-prey oscillators constitutive sequences**

#### 1.2. Reaction assembly

The buffer used for all experiments was composed of: 20 mM Tris-HCl (pH = 8), 10 mM (NH4)2SO4, 10 mM KCl, 8.4 mM MgSO4, 50 mM NaCl, 3 mM dithiothreitol (DTT), 2 µM Netropsin (Sigma-Aldrich), 400 µg.ml−1 BSA (New England Biolabs, NEB), 0.1 % Synperonic F108 (Sigma-Aldrich), 1x EvaGreen dye (Biotium), 100 nM Dextran Alexa Fluor 594 (Thermofisher) and dNTPs (200 µM each). Apart from predation and degradation characterization experiments (Figures S1 & S2), polymerase, nickase and exonuclease were added to the mix. The nicking endonuclease Nb.BsmI (NEB) was used at 400 U.mL−1 (4% of commercial stock solution). Bst DNA Polymerase, Full Length (NEB) was used in a range varying from 50 to 75 U.mL−1 (1 % to 1.5 % of commercial stock solution). ETSSB (NEB) was used in a range varying from 5 to 20 µg.mL−1 (0.5 % to 1.5 % of commercial stock solution). The thermophilic 5′–>3′ exonuclease from Thermus thermophilus ttRecJ was expressed and purified in-house as previously described [2]. The stock solution of ttRecJ was diluted to 1.53 µM in Diluent A (NEB) supplemented with 1% Triton X-100, and stored at −20 °C. ttRecJ was used in a range varying from 15.3 nM to 45.9 nM (1 % to 3 % of the dilution).

.

Concentrations for each figure are listed below:

- **Figure 1 D,F:** G1 60 nM, N1 1 nM, P1 9 nM, Polymerase 62.5 $U.mL^{-1}$, Nickase 400 $U.mL^{-1}$, Exonuclease 29 nM, ETSSB 12.5 $\mu g.mL^{-1}$
- **Figure 1 E:** (see SI 2.2)
- **Figure 4 :** G1 60 nM, N1 1 nM, P1 9 nM, Polymerase 62.5 $U.ml^{-1}$, Nickase 400 $U.ml^{-1}$, Exonuclease 29 nM, ETSSB 12.5 $\mu g.mL^{-1}$ (same as Figure 1D)
- **Figure 5:** G1 50 nM, N1 6 nM, P1 4 nM, Polymerase 75 $U.mL^{-1}$, Nickase 400 $U.mL^{-1}$, Exonuclease 21 to 30 nM, ETSSB 10. $\mu g.mL^{-1}$
- **Figure 6 :** G1 60 nM, N1 1 nM, P1 9 nM, Polymerase 62.5 $U.mL^{-1}$, Nickase 400 $U.mL^{-1}$, Exonuclease ranging from 15.3 nM to 45.9 nM, ETSSB 12.5 $\mu g.mL^{-1}$

All reaction components were assembled on ice to prevent the early start of the reaction. Bulk experiments were performed in a total volume of 8 µL and run in a CFX96 real-time PCR detection system (Biorad). Chemical neurons presented in Figure 6 were encapsulated into droplets of varying concentrations of exonuclease following previously published protocols [3,4] and incubated in a Silicon chamber placed in a temperature gradient[5].

#### 1.3. Droplet microfluidics methods

In short, we used a flow focusing PDMS chip to create a mono-disperse water in oil emulsion of ~50 µm droplets. The PDMS slab was obtained via soft-lithography from a SU8 mold (height ~55µm) and then bonded to a ~1mm thick glass slide using $O_2$ plasma. In the chip two aqueous channels containing respectively a high and low concentration of exonuclease merge and intersect an oil channel containing fluorinated oil (HFE 7500, Novec) and 3% surfactant (Emulseo, France), which triggers the formation of droplets at the exit of a nozzle. We mapped exonuclease concentration by precisely exchanging the flows of high and low exonuclease concentration channels using a pressure controller (MFCS-EZ from Fluigent, France). The emulsion was collected at the outlet in a pipette tip.

We filled a 3 cm x 1cm x 55µm silicon chamber with the emulsion to form a monolayer of droplets and closed it with a 165µm thick coverslip. The silicon chambers were fabricated in the clean room of the FEMTO-ST lab (Besancon, France), using standard lithography methods and Deep Reactive Ion Etching. After reception of the wafer, we spin-coated with a 10% Cytop CTL-809M (Asahi Glass) on the chambers and coverslips to render them hydrophobic and stabilize the droplets.

To incubate the droplets in a temperature gradient, the chamber was placed at the center of a copper-plate (16 cm x 4 cm x 0.5 cm). To impose a gradient of 5°C/cm, we used two Peltiers elements (Adaptive, 40 x 40 mm ET-161-12-08-E) paired with a controller (TEC-1122, Meerstetter) and Pt-100 sensors (RS-Pro, 10 mm x 2 mm probe, 4-wire, Class A). The error on the gradient is in the 0.1°C/cm range. To make comparison possible between different experimental setup, we harmonized temperature between droplets and bulk experiments. To do that, we used the temperature that maximizes the oscillation frequency (Figure 2D) as the reference temperature in droplets (which amounts to offsetting the temperature) .

We imaged the chamber using a motorized Nikon Ti2-E epifluorescence microscope connected to a LED light source (pE-4000, CoolLed) and a sCMOS camera (Prime 95B 25 mm, Photometrics). We used a 10x objective (CFI Plan Apo Lambda S10X, NA=0.45, Nikon) and approriate filters (purchased from Semrock or Chroma). Before analysis, images were unshaded using BaSiC[6] plugin and stitched on ImageJ[7].

Microscopy images were analysed using Mathematica following previously described protocols [3,4]. Briefly droplets were detected using the dy530 channel. The temperature inside the droplets was linearly interpolated from their position with respect to the two Peltier Element (which fixed the boundary conditions). The concentration of the exonuclease was inferred from the fluorescence of its dextran barcode. Once the fluorescence levels of each droplet is recovered the rest of the analysis is similar to bulk experiments.

#### 1.4. Fluorescence data analysis

Predator and prey populations are monitored in real-time using fluorescence. When a prey binds to the grass template it quenches the dy530 template's dye (Figure 1C). This N quenching is reversible and proportional to the concentration of prey. Therefore, negative peaks of yellow fluorescence indicates a surge and a fall in the population of prey. In addition, the EvaGreen dye added to the solution intercalates in DNA, giving a non-specific signal that is proportional to the total population of prey and predator (it gives a stronger weight to predators due to their hairpin secondary structure). The Dextran Alexa Fluor 594 (at a constant concentration) is used as a reference signal to check for errors of volume or evaporation. Fluorescence was acquired every 2 minutes in each of the 3 channels.

For clarity, after baselines removal, all dy530 signals were rescaled by the same factor in a given experiment. In Figure 4 the signals corresponding to a hot temperature were offset to account for the small variation of dy530 fluorescence (~15 % for 10°C). To account for the variation of EvaGreen fluorescence with temperature, EvaGreen fluorescence signals in Figures S2 and S3, were normalized respectively by their final or initial value corresponding to a known concentration of predators or preys (all preys transformed into predators for Figure S2 and all DNA strands degraded for Figure S3.

In Figure 4, for each tube, for each step of 2 hours we consider the output a "1" if there is a least spike of prey concentration, "0" if not. The consensus output corresponds to the bit of at least two of the triplicates.

In Figure 1F and Figure 6D, frequency was obtained by detecting the peaks in the signal using Mathematica, extracting all the times between consecutive peaks and taking the inverse of the average of these times. In Figure 6D, droplets with less than 3 peaks were considered "poor oscillators" and not considered for this calculation but used to define the "poor oscillations" zone appearing on the diagram.

### 2. Chemical Neuron Toy model

#### 2.1. Predator Prey equations

We mathematically describe the dynamics of predators and preys with the model of [1], to which we added temperature variations that we extracted experimentally, and a very small positive term $\varepsilon$ in each equation to account for the spontaneous emergence of preys or predators. For simplicity's sake, we assume that $\varepsilon$ does not depend on the temperature. The following equations are referred to as the governing equations for preys and predators in the following study.

$$\begin{cases} \dot{x}(t) = A(T) \cdot \dfrac{g \cdot x(t)}{1 + \beta \cdot g \cdot x(t)} - B(T) \cdot y(t) \cdot x(t) - C_x(T) \cdot \dfrac{\lambda \cdot \delta \cdot x(t)}{1 + y(t)} + \varepsilon \\ \dot{y}(t) = B(T) \cdot y(t) \cdot x(t) - C_y(T) \cdot \dfrac{\delta \cdot y(t)}{1 + y(t)} + \varepsilon \end{cases}$$

where t is the normalized time (scaled by $t_c$ ) $x$, $y$ and $g$ are the normalized concentrations of respectively prey, predator (scaled by the Michaelis Menten constant of the exonuclease $K_m$) and template. $\beta$ is a saturation constant. $\delta$ and $\lambda \cdot \delta$ are the normalized decay rates of predators and preys respectively. $A$, $B$, $C_x$ and $C_y$ denote the temperature dependence of prey replication, predation, prey degradation, predator degradation respectively. Each one of these prefactors is equal to 1 at 46.5°C, so that we recover the equation [1] at the temperature they used. We fixed the following parameters: $K_m = 35$ nM, $\beta = 0.08$, $g = 2.3$, $\delta = 0.17$, $t_c =$ 2.3 min, $\lambda = 5$. $K_m$ and $\beta$ were directly taken from [1]. $\lambda$ was extracted from exonuclease decay rates experiment (see section 2.2.3). $t_c = 2.3$ min was extracted to adjust the period of simulated oscillation with experimental data from Figure 1.

The values of $g$ and $\delta$ were adjusted to account for the change in activity between our polymerase and the one used 8 years ago by Fujii and Rondelez. Indeed, using the experimental traces from Figure S11 in the supplementary materials of Fujii & Rondelez, 2013 [1] which corresponds to a similar concentration of exonuclease as our experiments, we can assess that with our polymerase we obtain similar oscillations with 60 nM of template than those obtained for ~120 nM in S11. This means that the activity of our polymerase is double. Accordingly, $\delta$ being the ratio of the activity of the exonuclease normalized by the activity of the polymerase we divided it by a factor of 2.

We chose $\varepsilon = 10^{-8}$ to account for the self-start of PEN Toolbox amplification reactions (i.e. the emergence of predator and prey from ab-initio synthesis by the polymerase). This mechanism acts as a constant but small source of predator and preys. Experimentally, it is difficult to separate from a contamination of predators or preys in the fM range [8]. This parameter $\varepsilon$ is also helpful to numerically stabilizes the simulations. It prevents concentrations from reaching exponentially low levels (where errors from floating points computations can easily corrupt computations).

#### 2.2. Extraction of the temperature dependence of the prefactors

##### 2.2.1. Prey replication : A(T)

We experimentally studied the variation of amplification times with temperature for different initial concentrations of prey. To do so, we isolated the prey replication block by removing the initial predators and the exonuclease, and put an excess of dNTPs to consider that enzymatic production is constant over time. Those conditions allow us to only consider the replication term in the prey governing equation:

$$\dot{x}(t) = A(T) \cdot \frac{g \cdot x(t)}{1 + \beta \cdot g \cdot x(t)}$$

which is integrated into

$$\beta \cdot \big(x(t) - x(0)\big) + \frac{1}{g} \cdot \ln\left(\frac{x(t)}{x(0)}\right) = A(T) \cdot t$$

For any threshold value $x(t_s) = x_s \gg x(0)$, we can derive the time over threshold, $t_s$ :

$$t_s(T) = \frac{1}{A(T)}\left(\beta \cdot x_s + \frac{1}{g}\ln(x_s)\right) - \frac{1}{g \cdot A(T)} \cdot \ln\big(x(0)\big)$$

In practice, that threshold value is set to correspond to a near maximum fluorescence value. For a given temperature, we obtain $\frac{1}{g \cdot A(T)}$ by linear regression of $t_s$ against $-\ln\big(x(0)\big)$. Experimental data for times over threshold fits well to a linear evolution (Fig S1 C). To account very simply for the non-monotony of $g \cdot A(T)$ we fit it with a quadratic function and obtain A(T) by normalizing at 46.5 °C (Fig S1 D).

$$A(T) = a \cdot T^2 + b \cdot T + c$$

with $a = -8.2 \cdot 10^{-3}\ °C^{-2}$, $b = 0.76\ °C^{-1}$ and $c = -16.7$

The non-monotony can be explained by the balance between input hybridization on the template which is favored at lower temperatures and enzyme activity which increases with temperature.

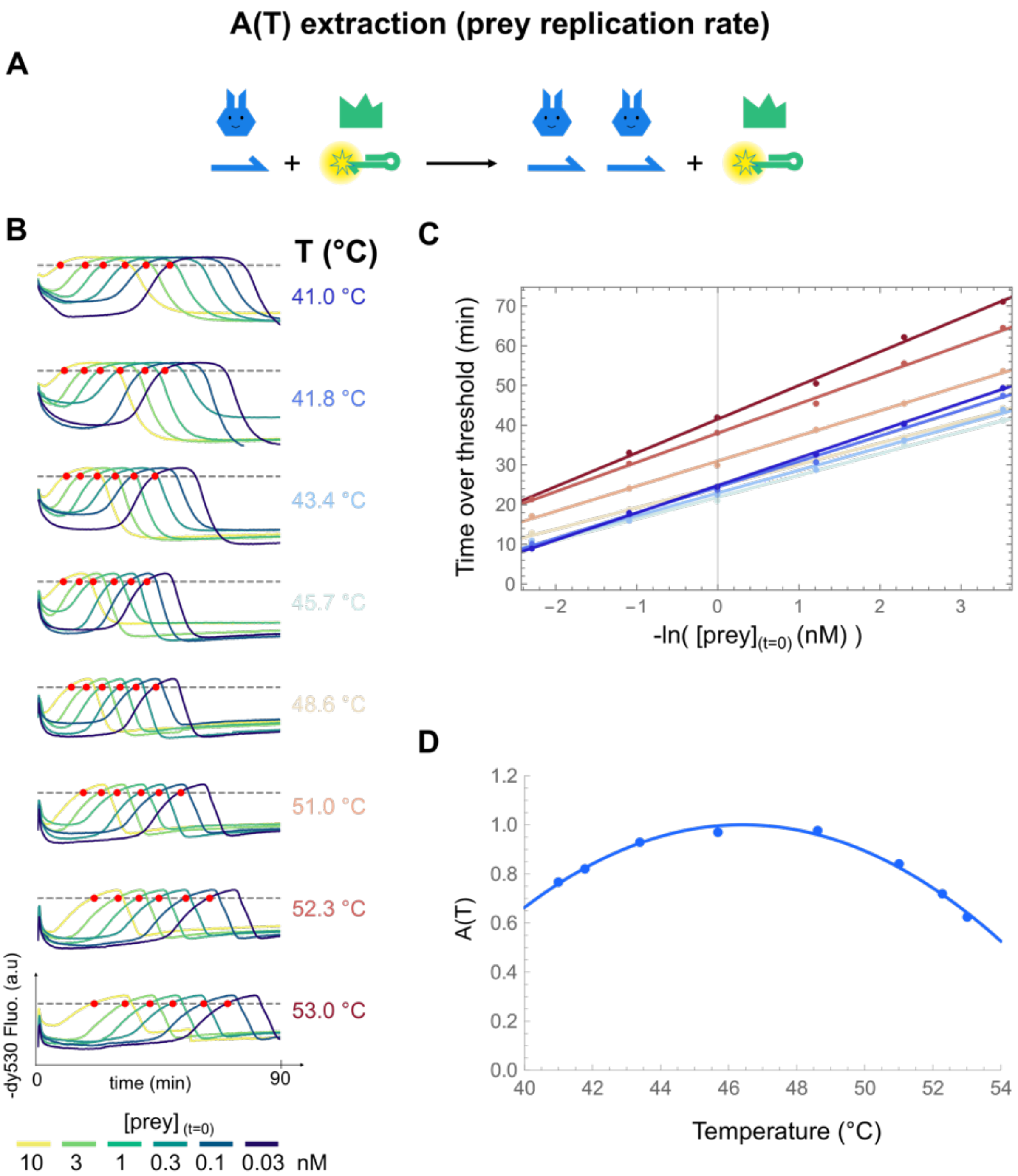

**Figure S1. Prey replication kinetics temperature dependence.** (**A**) Prey replication reaction. (**B**) Prey amplification curves for different initial prey concentrations at different temperatures. Red dots note the passage of the threshold. (**C**) Linear fit of the experimental times over threshold against the natural logarithm of prey concentration. (**D**) Polynomial fit of the variation of coefficient A with temperature.

##### 2.2.2. Predation: B(T)

Having removed the G1 template and the exonuclease to isolate the predation pathway (Fig. S2 A), 100 nM of prey and 10 nM of predators were transformed into 110 nM of predators for temperatures ranging from 40 °C to 52 °C. The transformation of preys into predators is monitored via EvaGreen fluorescence which is higher for predators than for preys due to their hairpin structure (Fig S2 B)

In absence of exonuclease, and template we only consider the predation term in the predator and prey equations:

$$\begin{cases} \dot{x}(t) = -B(T) \cdot y(t) \cdot x(t) \\ \dot{y}(t) = B(T) \cdot y(t) \cdot x(t) \end{cases}$$

The sum of predators and preys remains constant over time and we can write

$$\dot{y}(t) = B(T) \cdot y(t) \cdot \left( x(0) + y(0) - y(t) \right)$$

Which by integration gives:

$$\frac{1}{\mathrm{x}(0) + y(0)} \left( \ln\left( \frac{y(t)}{y(0)} \right) - \ln\left( \frac{x(0) + y(0) - y(t)}{x(0)} \right) \right) = B(T) \cdot t$$

By placing a threshold when half of the preys are consumed, $y(t_b) = y(0) + \frac{x(0)}{2}$, we obtain times over threshold (Fig. S2 C) inversely proportional to B(T). By plotting the inverse of those times and normalizing at 46.5 °C we obtain B(T). We fit B(T) with a sigmoid function (Fig S2 D).

$$\mathrm{B(T)} = \frac{\mathrm{a}}{1 + \exp\left(-b \cdot (T - T_s)\right)}$$

with $\mathrm{a} = 2$, $\mathrm{b} = 0.33\ °C^{-1}$, $T_s = 46.5\ °C$. Predation is nearly 5 times faster at 52°C than at 42°C. This increased rate and its sigmoidal shape can be explained by the dehybridization of the newly formed predators at high temperature which allows them to capture new preys. The sigmoid is not centered on the pure melting temperature of two predators, which is 54°C [1], because prey hybridization on the predator and strand displacement effect of the BST polymerase shift the predation sigmoid to lower temperatures.

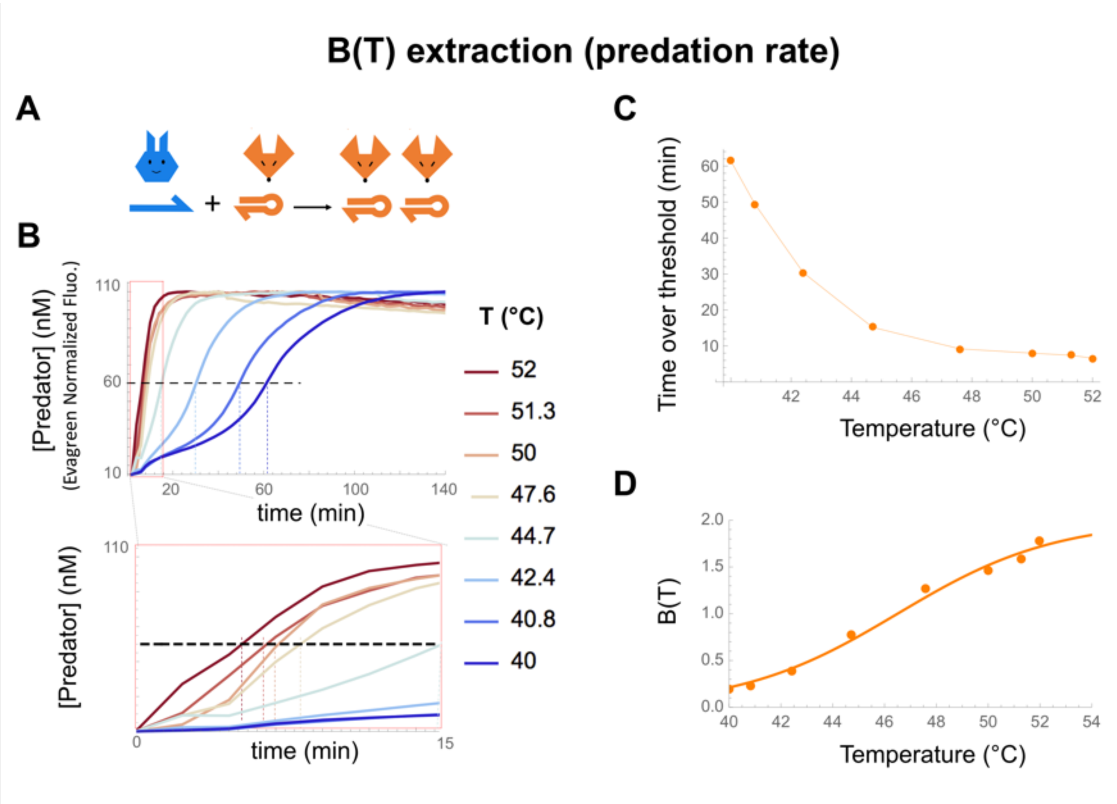

**Figure S2. Predation kinetics temperature dependence. (A)** Predation reaction. **(B)** Evolution of the concentration of predators over time at different temperatures. (**C**) Times over threshold plotted against temperature. (**D**) Sigmoidal fit of the variation of B(T). N1 100 nM, P1 10nM, Polymerase 62.5 U.ml$^{-1}$, Nickase 400 U. ml$^{-1}$, ETSSB 10 µg µg.mL$^{-1}$, No Template G1, and no exonuclease were used to observe only predation.

##### 2.2.3. Degradation: Cx(T) and Cy(T)

To measure degradation rates, we incubated preys or predators with exonuclease only. We used large initial concentrations of preys or predators ($c_0 = 2$ µM) to saturate the exonuclease and measured the decay rates of each species. The governing equations become:

$$\begin{cases} \dot{x}(t) = -C_x(T) \cdot \lambda \cdot \delta \\ \dot{y}(t) = -C_y(T) \cdot \delta \end{cases}$$

by integration we get:

$$x(t) = c_0 - C_x(T) \cdot \lambda \cdot \delta \cdot t$$

$$y(t) = c_0 - C_y(T) \cdot \delta \cdot t$$

By taking a concentration threshold at $c_s = 0.6\, c_0$ we obtain times over threshold $t_{sx}$ and $t_{sy}$ inversely proportional to $C_x(T)$ or $C_y(T)$ respectively:

$$\frac{1}{t_{sx}(T)} = C_x(T) \cdot \lambda \cdot \frac{\delta}{c_0 - c_s}$$

$$\frac{1}{t_{sy}(T)} = C_y(T) \cdot \frac{\delta}{c_0 - c_s}$$

We fit $C_x(T)$ and $C_y(T)$ with an Arrhenius law to model the thermophilicity of the ttRecJ exonuclease. Indeed, this protein is extracted from Thermus thermophilus bacterium whose optimal growth occurs between 65°C and 72 °C [9]

$$C_x(T) = a \cdot \exp\left(-\frac{E_x}{R(T - T_0)}\right)$$

$$C_y(T) = b \cdot \exp\left(-\frac{E_y}{R(T - T_0)}\right)$$

where $T_0 = -273{,}15$ °C is the absolute 0 temperature and $R = 8.314\, \mathrm{J \cdot °C^{-1} \cdot mol^{-1}}$ the universal gas constant. $E_x$ and $E_x$ are activation energies that can be obtained by plotting the inverse of the time over threshold against the inverse of temperature (Fig. S3 D):

$$\ln\left(\frac{1}{t_{sx}(T)}\right) = -\frac{E_x}{R(T - T_0)} + \ln(a) + \ln\left(\lambda \cdot \frac{\delta}{c_0 - c_s}\right)$$

$$\ln\left(\frac{1}{t_{sy}(T)}\right) = -\frac{E_y}{R(T - T_0)} + \ln(b) + \ln\left(\frac{\delta}{c_0 - c_s}\right)$$

The linear fit gives $E_x = 1.43 \cdot 10^5$ J $\cdot$ mol$^{-1}$ and $E_y = 1.83 \cdot 10^5$ J $\cdot$ mol$^{-1}$, while normalizing at 46.5 °C gives $a = 1.62 \cdot 10^{24}$ and $b = 1.25 \cdot 10^{30}$. Before normalization we can also extract $\lambda = 5$ which is in agreement with the value $\lambda = 4.5$ that we find in the literature [1].

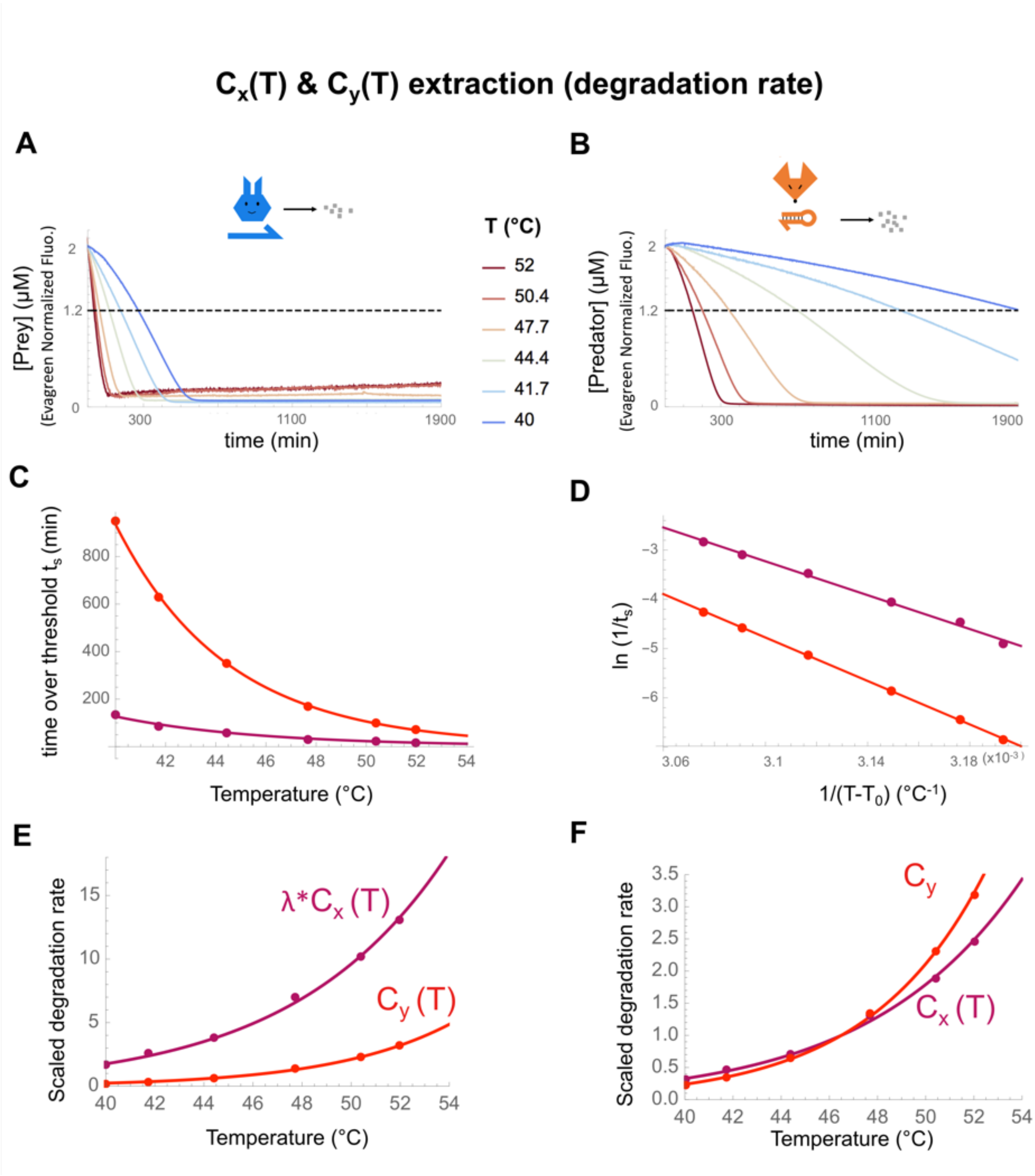

**Figure S3 - Characterization of the evolution of the degradation of prey and predator strands with temperature.** (**A**) Degradation of preys monitored with EvaGreen. (**B**) Degradation of predators monitored with EvaGreen. (**C**) Times over threshold plotted against temperature. (**D**) Extraction of activation energies. (**E**) Degradation speed of preys and predators. (**F**) Normalized degradation speed of preys and predators. N1 2 or 0 µM , P1 0 or 2 µM, Exonuclease 30.6 nM, ETSSB 10 µg.mL−1. No Template G1, no polymerase and no nickase were used to observe only degradation by exonuclease.

#### 2.3. Evolution of the Frequency with temperature

To apprehend the effect of temperature on the frequency of oscillations, we linearized our system, reducing it to the canonical Lotka-Volterra oscillator. This well-studied system admits an analytical derivation of the frequency of its oscillation[10] – a derivation that is based on a linear approximation used by both Lotka [11] and Volterra [12].

$$\begin{cases} \dot{x}(t) = x(t) \cdot (a - b \cdot y(t)) \\ \dot{y}(t) = y(t) \cdot (c \cdot x(t) - d) \end{cases}$$

with $a = A(T) \cdot g - C_x(T) \cdot \lambda \cdot \delta,\ b = c = B(T),\ d = C_y(T) \cdot \delta$

This system has only one fixed point $(x_f, y_f) = (\frac{d}{c}, \frac{a}{b})$ around which periodic solutions oscillate. The linearized system around this point is:

$$\begin{cases} \dot{x}(t) = x_f \cdot (a - b \cdot y(t)) \\ \dot{y}(t) = y_f \cdot (c \cdot x(t) - d) \end{cases}$$

For strictly positive initial conditions $(x_0, y_0)$ the system has the following periodic elliptic solutions:

$$x(t) = \frac{d}{c} + r \cdot \sin\,(t \cdot 2\pi \cdot f + t_*)$$
$$y(t) = \frac{a}{b} + r \cdot \frac{c}{b}\sqrt{\frac{a}{d}} \cos\,(t \cdot 2\pi \cdot f + t_*)$$

with $f = \frac{\sqrt{ad}}{2\pi}$ the frequency and $r = \sqrt{\left(x_0 - \frac{d}{c}\right)^2 + \left(y_0 - \frac{a}{b}\right)^2 \frac{b^2 d}{ac^2}}$ the amplitude of the oscillations. As $b = c$ and considering $x_0 \ll 1$ and $y_0 \ll 1$ we can get $r$ as

$$r = \frac{d}{b}\sqrt{1 + \frac{a}{d}}$$

which for our system corresponds to:

$$r = \frac{C_y(T) \cdot \delta}{B(T)} \sqrt{1 + \frac{A(T) \cdot g - C_x(T) \cdot \lambda \cdot \delta}{C_y(T) \cdot \delta}}$$

This expression shows that strong predation $B(T)$, decreases the amplitude $r$ while the net prey growth $A(T) \cdot g - C_x(T) \cdot \lambda \cdot \delta$ increases it which agrees with intuition. The frequency for the linearized system is:

$$f = \frac{\sqrt{(A(T) \cdot g - C_x(T) \cdot \lambda \cdot \delta) \cdot C_y(T) \cdot \delta}}{2\pi}$$

If $A(T) \cdot g - C_x(T) \cdot \lambda \cdot \delta = 0$, the system undergoes a bifurcation where the frequency reaches 0 and oscillations stop. For our system this happens for T ~ 51°C (Fig. 1F) which is within ~1°C of the experimental value (Fig. 1D).

The oscillation frequency in the linearized model is the geometrical mean of the net growth of prey and the decay of predators. Indeed, one oscillation is mainly composed of two phases: first a prey growth which is quasi exponential, and then an exponential decay of predators once they have depleted the preys. High frequencies come from a fast growth of preys or a fast decay of predators.

This derivation explains the non-monotony of frequency with temperature as degradation – the pathway that is the most sensitive to temperature (Fig. 1E) – appears in both terms of the geometrical mean. At low temperatures $A(T) \cdot g \gg C_x(T) \cdot \lambda \cdot \delta$ therefore temperature mostly affects the predator decay $C_y(T) \cdot \delta$ and an increase in temperature increases frequency. Once $A(T) \cdot g \sim C_x(T) \cdot \lambda \cdot \delta$ increasing temperature drastically delays prey growth which translates in a drop of frequency (Fig. 1F).

#### 2.4. Prediction of bifurcation temperature

Sustained oscillations require predators and preys, and a state with only preys should be unstable and lead to the growth of predators. At a given temperature, we consider first the equilibrium for preys in absence of predator and then study the resulting net growth of predators.

In the absence of predators and neglecting the $\varepsilon$ term, the governing equation for preys becomes at steady state:

$$0 = (A(T) \cdot \frac{g}{1 + \beta \cdot g \cdot x_s} - C_x(T) \cdot \lambda \cdot \delta) \cdot x_s$$

which gives for the steady state concentration of preys

$$x_s = \frac{1}{\beta} \cdot \left( \frac{A(T)}{C_x(T)} \cdot \frac{1}{\lambda \cdot \delta} - \frac{1}{g} \right)$$

The governing equation for predators, considering $y(t) \ll 1$, becomes:

$$\dot{y}(t) = (B(T) \cdot x_s - C_y(T) \cdot \delta) \cdot y(t) + \varepsilon$$

To obtain a net growth of predators (Figure 1G) that is significant (and not in the $\varepsilon$ range), we must have:

$$B(T) \cdot x_s - C_y(T) \cdot \delta > 0$$

That condition makes intuitive sense, as it means that predator growth occurs when the consumption of prey outpaces the death of predators.

#### 2.5. Numerical study of the nature of the different steady states and bifurcation of the chemical neuron

Local bifurcations correspond to the appearance/disappearance of a steady state, or a change in its stability. To track local bifurcations, we study the Jacobian matrix J of our governing equations,

$$J = \begin{pmatrix} \frac{\partial \dot{x}}{\partial x} & \frac{\partial \dot{x}}{\partial y} \\ \frac{\partial \dot{y}}{\partial x} & \frac{\partial \dot{y}}{\partial y} \end{pmatrix}$$

We change temperature, and track how the eigenvalues of J, evaluated at each steady state, vary with temperature. The nature of the eigenvalues of J reflects the local dynamic around the steady state. Eigenvalues with a non-null imaginary part reflects an oscillatory behavior. Eigenvalues with a real part that is negative locally stabilize the system (attracting the trajectories to the steady state), while eigenvalues with a real part that is positive locally destabilize the system (expelling the trajectories from the steady state).

Note that we do not actually need to numerically find the eigenvalues of the Jacobian. For a two-dimensional dynamic system, the two eigenvalues of J are entirely determined by the trace and the determinant of J, which are the sum and product of the eigenvalues respectively. In other words, the qualitative nature of a steady state at a given temperature can be visualized from the quadrant to which belong the point (tr(J), det(J)). By plotting the trajectories followed by these points as the temperature parameter is varied, we can visualize the bifurcation route of the system (Figure S4).

There are 3 steady states (not always co-existing at the same time): a stable node, a saddle and a spiral. The 3 distinct trajectories merge into one line, but the saddle point travels in the opposite direction and collides with the two other states. The collision occurs on the tr(J) axis which attests of saddle node bifurcations. When the spiral state crosses the det(J) axis, a Hopf bifurcation occurs: its stability switches. This creates a limit cycle.

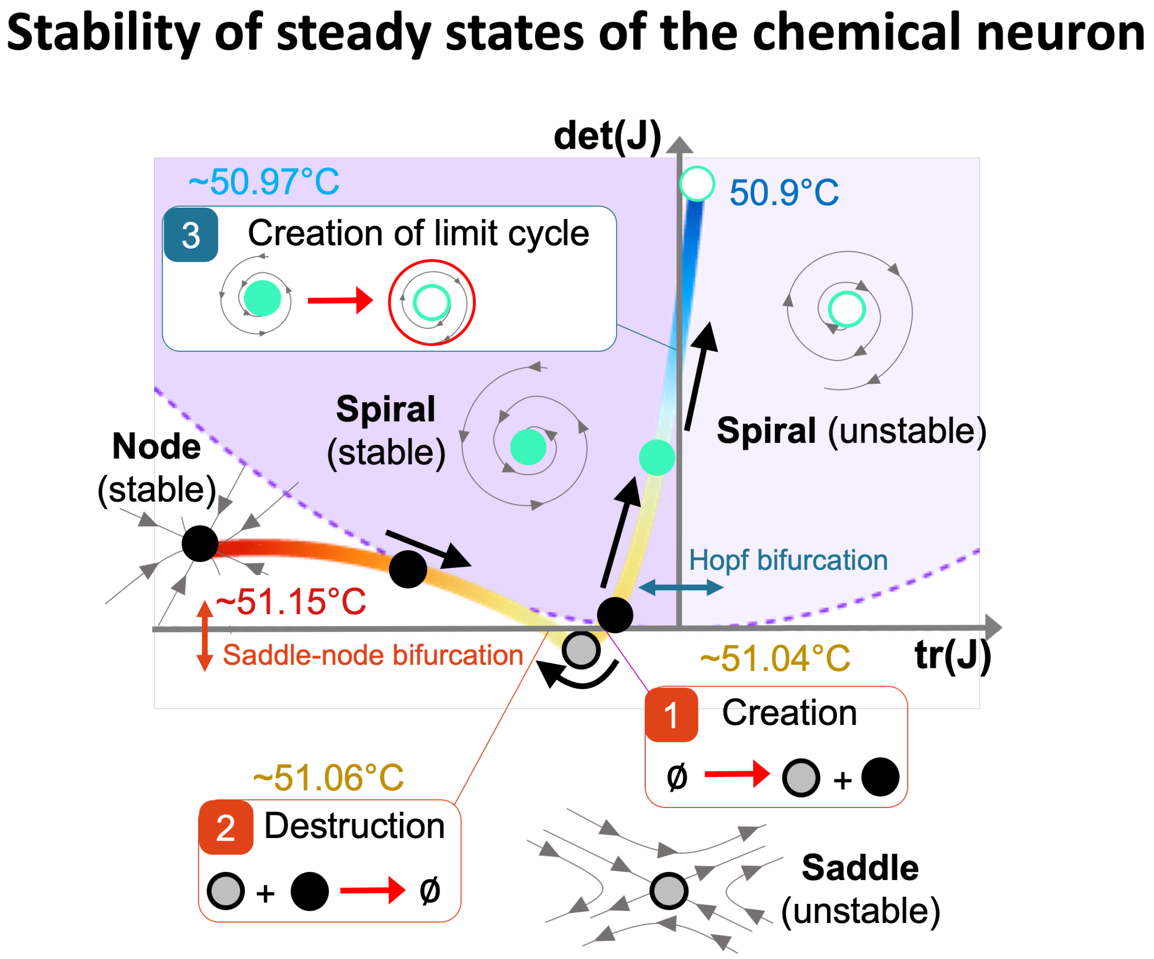

**Figure S4: (Trace,Determinant) diagram for the Jacobian matrix J of the chemical neuron at each stady state.** The zone above the purple dotted parabola corresponds to eigenvalues with an imaginary part. A steady state crossing the det(J) axis correspond to a Hopf bifurcation while two points colliding on the tr(J) axis correspond to a saddle-node bifurcation. The trajectories of the 3 steady states (not always existing at the same time) are color coded by the temperature.

#### 2.6. Bifurcation diagram

We experimentally mapped the 2D bifurcation diagram of the neuron against temperature and exonuclease (Fig. S5A), and confronted it to the predictions of the model (Fig. S5B). Overall the bifurcation temperature decreases with increasing exonuclease - a dependence that is correctly predicted by the model, and tracked with a precision of a few degrees °C.

Obviously, the model does not account for enzymatic fatigue, that is the loss of enzymatic activity with time, which likely explains the shape of oscillations at the highest concentration of exonuclease and the lowest temperature (bottom right in the diagrams). There, the amplitude of oscillations decreases and then increases, a non-monotonic behavior that is not captured by the model.

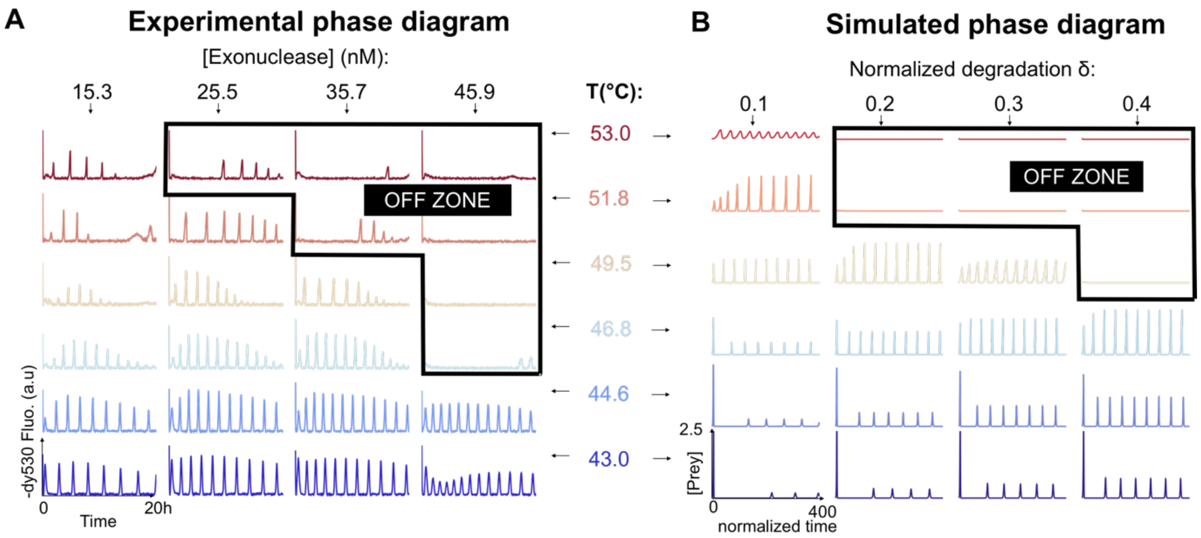

**Figure S5. Phase diagrams in the plane (Exonuclease, Temperature) of the chemical neurons** (**A**) Experimental phase diagram in bulk. Tuning the concentration of exonuclease changes allows the temperature of bifurcation. Before being completely turned off at high temperatures, a time-lag - a signature of a saddle remnant – can be observed. (**B**) Predicted phase diagram. Both high temperature and high exonuclease concentration turn the oscillations off.

### 3. Biological thermal sensory neuron toy model

#### 3.1. Model description

For the sake of simplicity and for a direct analogy with our 2D predator prey toy model we chose a two-dimensional Hindmarsh–Rose type model as a toy-model of a thermosensory neuron[13]

$$\begin{cases} \dot{x}(t) = c\left(x(t) - \dfrac{x^3(t)}{3} - y(t) + I(T)\right) \\ \dot{y}(t) = \dfrac{x^2(t) + d\,x(t) - b\,y(t) + a}{c} \end{cases}$$

where $x$ denotes the cell membrane potential and $y$ a recovery variable. $I$ denotes the external stimuli, which for a thermosensory neuron depends of temperature T. Parametrization by $a$, $b$, $c$ and $d$ generates a large range of bifurcations, as extensively studied by [13]. Some bifurcations only occur in a region of the parameters space (Tsuji et al., 2007 Figure 5). We followed the bifurcation diagram presented in Tsuji et al., 2007 Figure 5, and set a=0.8, b=0.6, c=3, d=1.8. Such parameters beside giving similar nullclines to our chemical neuron also correspond to very large domains of similar behaviors.

To model the effect of a cold stimuli on our system causing the opening of TRPM8 channels, we modelled the external stimuli I by a sigmoidal current varying with temperature following previous experimental[14] and theoretical [15] studies.

$$I(T) = I_0\left(1 - \frac{1}{1 + \exp\left(-\frac{T - T_s}{\alpha}\right)}\right)$$

Where $I_0$ is the maximal normalized current, $T_s$ corresponds to the switching temperature where the current is half of $I_0$ and $\alpha$ gives the sharpness of the switch. We took $I_0 = 2$, $\alpha = 2\ °C^{-1}$, and $T_s = 25°\text{C}$. This gives an amplitude of variation of $0.85\ I_0$ between 20°C and 30°C (Fig. S4), which qualitatively agrees with experimental data [14]. This modelling of how temperature affect neurons is coarse-grained, and does not account for the transistor-like behavior of the TRPM8 channels. Unlike other studies that aimed at a better understanding of thermosensory neurons [15], our goal is simply to formulate a tractable toy model that captures salient experimental traits. The model exhibits firing/resting behaviors when submitted to a cold/hot stimuli, similar to what has been experimentally demonstrated on warm-inhibited afferents [16].

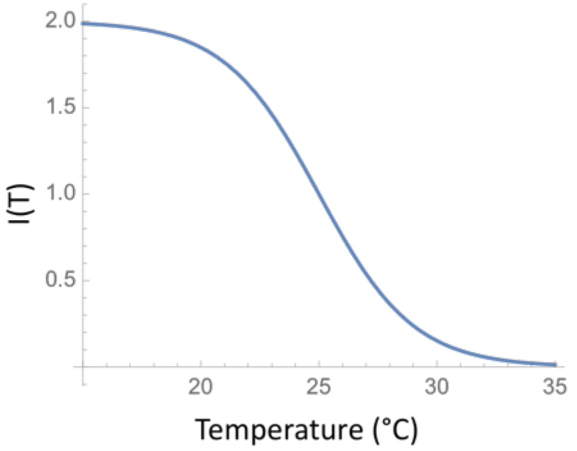

**Figure S6. External stimuli modelling the external current, resulting from the opening of TRPM8 channels at cold temperatures.**

#### 3.2. Numerical study of the nature of the different steady states and bifurcation of the thermal sensory neuron

Similarly to Figure S4, we numerically computed the steady states and the Jacobian matrix associated. When temperature is decreased a saddle and an unstable node are created. The creation of the limit cycle corresponds to the saddle exiting the oscillatory region. Further decreasing temperature destroys the stable resting state and allows for oscillations from any starting point (Figure S7).

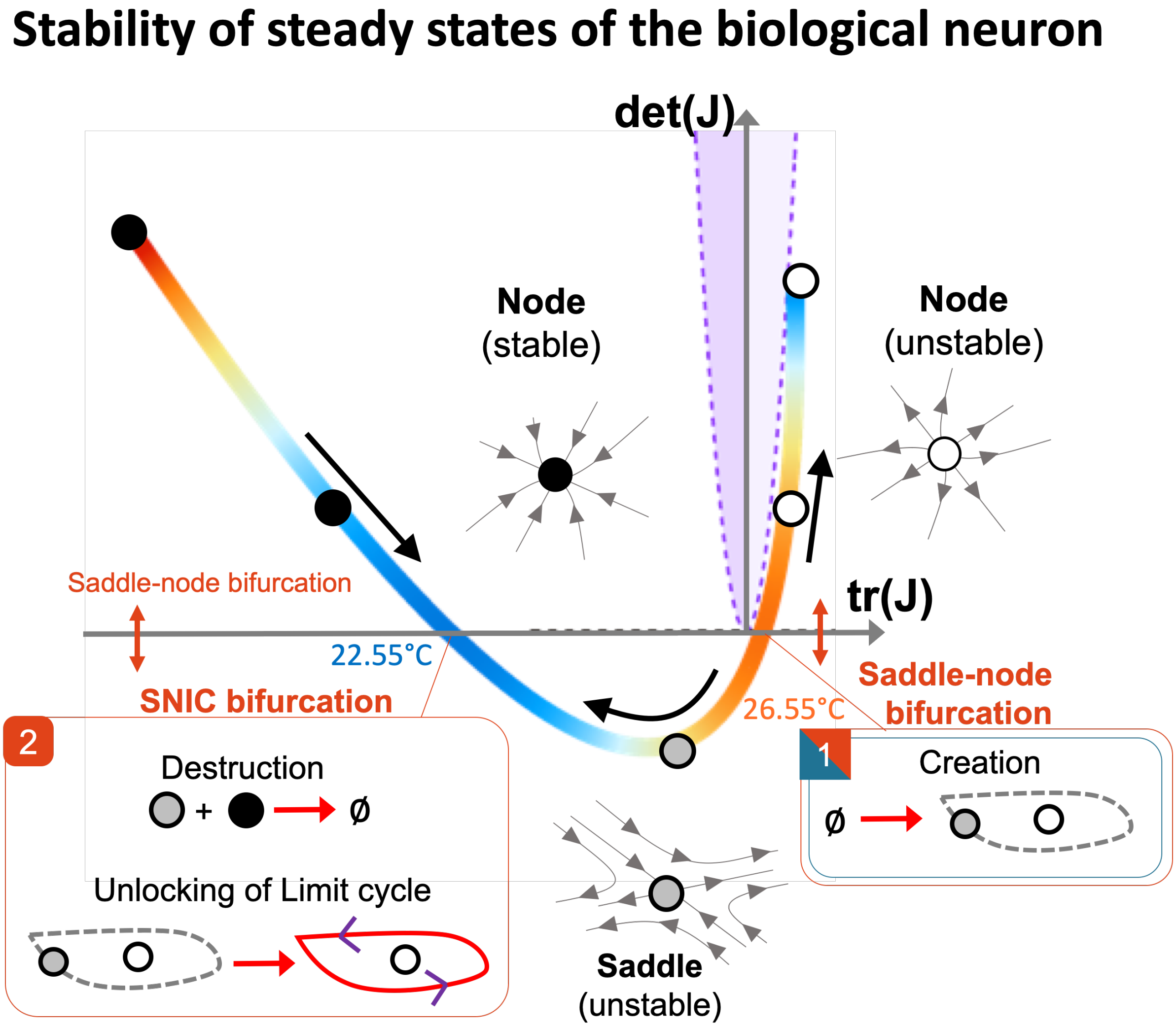

**Figure S7. (Trace,Determinant) diagram for the Jacobian matrix J of the biological neuron.** The trajectories of the 3 steady states (not always existing at the same time) are color coded by the temperature.

### 4. Canonical bifurcations between oscillations and rest

In this section, we qualitatively study canonical bifurcations between rest and oscillation.

#### 4.1. Homoclinic bifurcation

In an homoclinic bifurcation, a limit cycle grows until it collides with a neighboring saddle, which pries open the cycle and destroys oscillations (Figure S8).

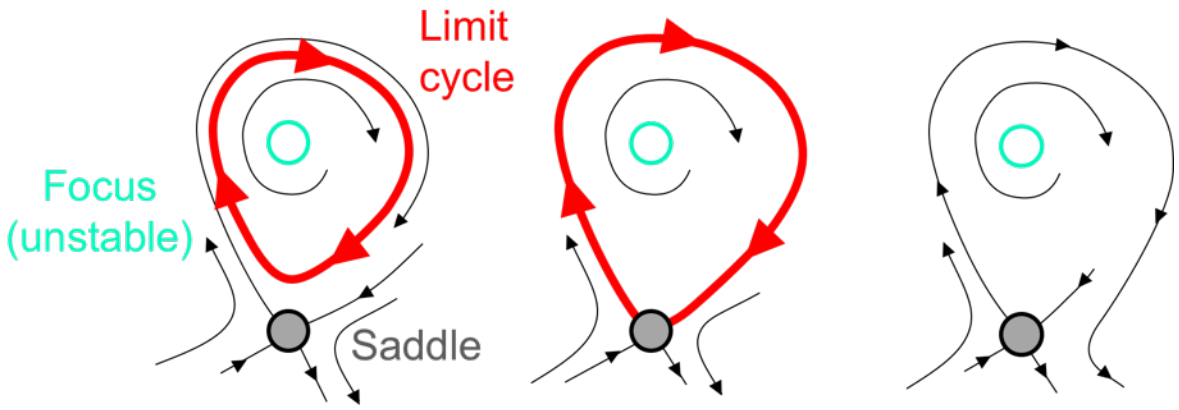

**Figure S8 Homoclinc bifurcation**

We used the following equations to model an homoclinic bifurcation parametrized by a bifurcation parameter $\mu$ (Fig. S9):

$$\begin{cases} \dot{x} = y \\ \dot{y} = \mu \cdot y + x - \dfrac{x^2}{1 + 0.01 \cdot x^2} + x \cdot y \end{cases}$$

In homoclinic bifurcations, when the control parameter is turned from ON to OFF, oscillations on a limit cycle are killed by the collision of a saddle node with the limit cycle - which opens the limit cycle, ejects the oscillating trajectory and switches the system from ON to OFF. But when the control parameter is switched back from OFF to ON, the ejected trajectory may not have remained close enough to the basin of attraction of the limit cycle to engage again in oscillations. Instead, the trajectory may have been trapped in the basin of attraction of the saddle node (which is never destroyed and still exists in the oscillation regime), or the trajectory may have escaped from the basins of the saddle node or the limit cycle altogether, and jumped to another region of the phase space (Figure S9). This manifests as the system remaining in an artifact OFF state (not oscillating) in spite of the control parameter being turned ON and OFF repeatedly (Fig. 3). This irreversibility is expected for a global bifurcation like the homoclinic bifurcation, because it alters globally the topology of the vector field, and does not guarantee that trajectories will remain confined during bifurcation.

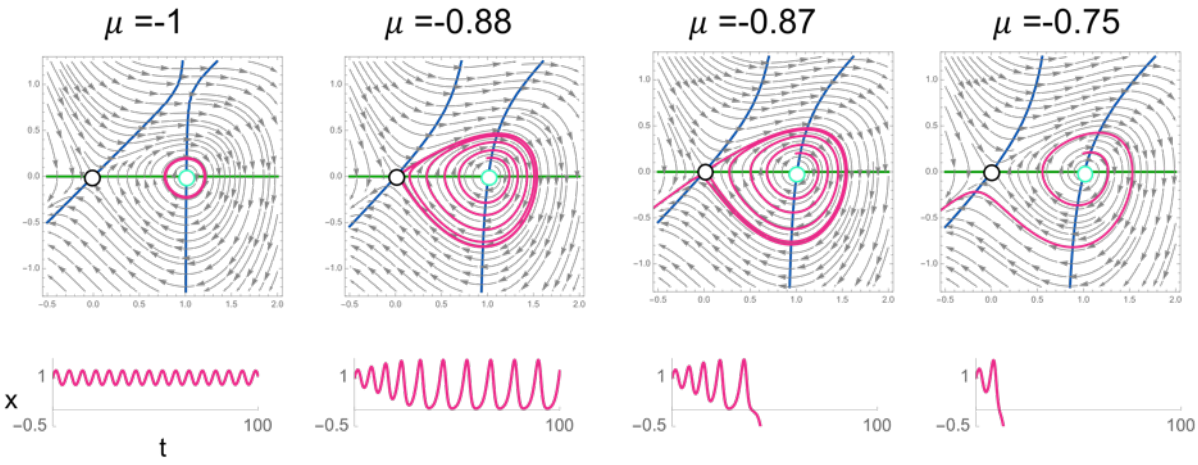

**Figure S9 Trajectories in a homoclinic bifurcation when $\mu$ is varied around the bifurcation point**

#### 4.2. Subcritical Hopf

In a subcritical Hopf bifurcation, a (stable) limit cycle shrinks around an unstable steady state, eventually disappearing into the steady state and making it stable (Figure S10)

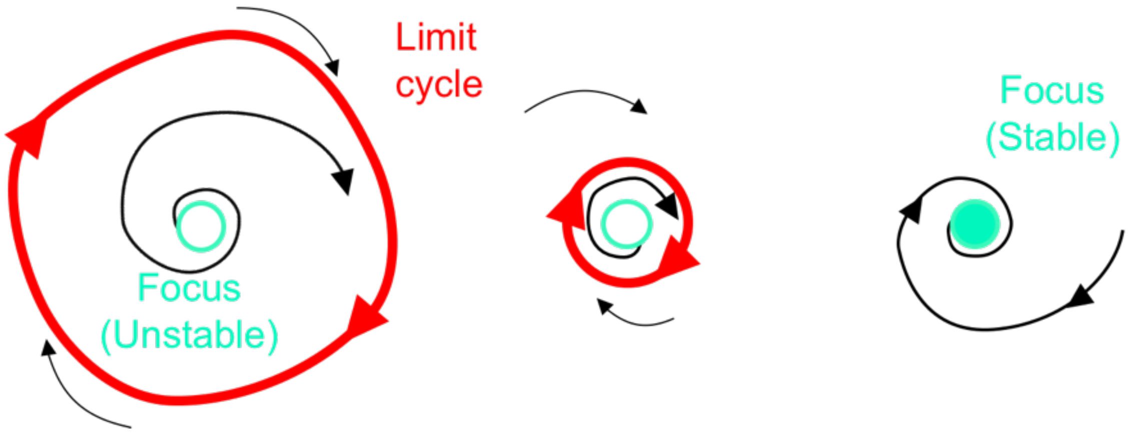

**Figure S10 Subcritical Hopf bifurcation**

We used the following equations to study the bifurcation when varying the bifurcation parameter $\mu$ (Fig. S11):

$$\begin{cases} \dot{x} = \mu \cdot x + y \\ \dot{y} = -x \cdot +\mu \cdot y - x^2 \cdot y \end{cases}$$

Consider a system with a supercritical Hopf bifurcation, with a control parameter μ that is ON but close to the bifurcation. The trajectory is oscillating on a limit cycle, but when the control parameter μ is turned OFF, the limit cycle shrinks and disappears into a spiral state (a steady state whose eigenvalues have a negative real part and a non-null imaginary part). The trajectory that was oscillating is now attracted to this steady state, which it reaches not by going in a straight line, but by "spiraling in" due to the presence of eigenvalues with imaginary values (which in a sense are the remnants of the limit cycle). This damping is obviously detrimental for spike-encoded sensing. When the stimulus is withdrawn (ON to OFF), the spiking sensor dampens until it stabilizes to the rest state. Conversely, when the stimuli is applied again, the sensor does not immediately produce spikes with correct amplitude, but goes through a transient regime where the amplitudes of the spike grows until they stabilize (Figure S8).

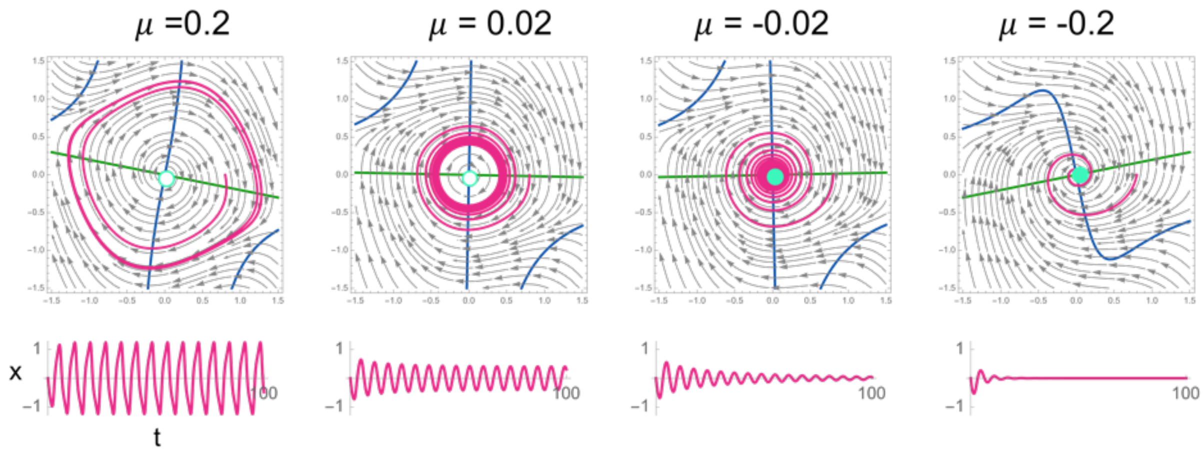

**Figure S11 Subcritical Hopf bifurcation when varying $\mu$ around the bifurcation point**.

#### 4.3. SNIC (Saddle-Node on an Invariant Cycle)/SNIPER bifurcation

In a SNIC bifurcation the limit cycle is never destroyed (it is invariant). Oscillations are stopped (started) by creating (destroying) a rest point on the limit cycle that traps the trajectory. Since the trajectory never leaves the limit cycle, oscillations are immediately restored to their fullest amplitude when the control parameter is switched back ON - avoiding the damping seen with the Hopf bifurcation. But the persistence of the limit cycle causes other artifacts, because the rest point only traps local trajectories on the limit cycle. When the system is turned from ON to OFF, oscillations that were sufficiently engaged past the rest point will continue their excursion along the limit cycle until they eventually meet the rest point - producing one erroneous spike. In other words, the system could spike once after withdrawal of the stimulus. In addition, the OFF state is sensitive to stimulus noise. Trajectory in the OFF state can escape the basin of attraction of the rest state thanks to transient noise, and engage in an excursion along the limit cycle - also producing an erroneous spike. This sensitivity to transient noise of SNIC bifurcation is sometimes desirable (for instance for neuronal excitability [17], but would be detrimental for a sensor which aims to faithfully represent the state of its environment.

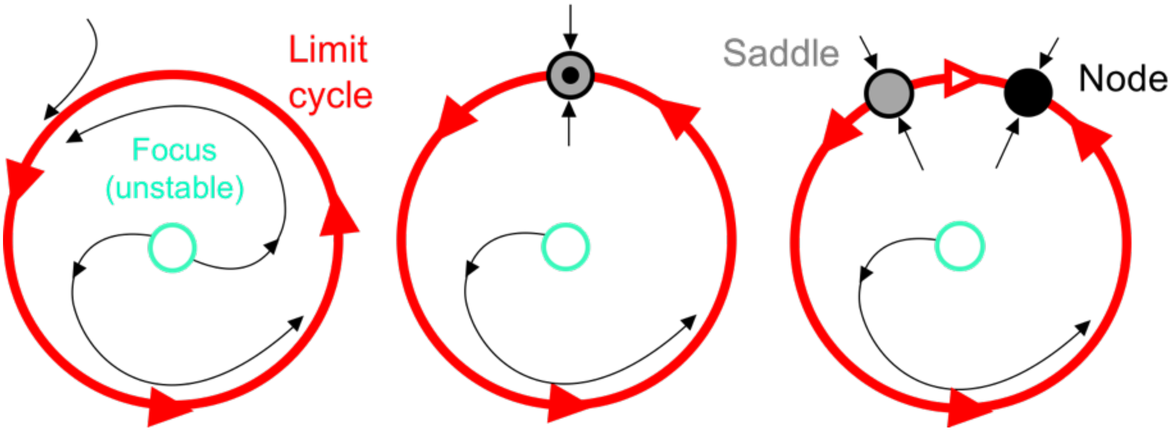

**Figure S12 A Saddle-Node on an Invariant Cycle bifurcation**

We used the following mode:

$$\begin{cases} \dot{r} = r \cdot (1 - r^2) \\ \dot{\theta} = \mu - \sin(\theta) \end{cases}$$

which in cartesian coordinates becomes:

$$\begin{cases} \dot{x} = x \cdot \left(1 - (x^2 + y^2)\right) - y \cdot (\mu - \frac{y}{\sqrt{x^2 + y^2}}) \\ \dot{y} = y \cdot \left(1 - (x^2 + y^2)\right) + x \cdot (\mu - \frac{y}{\sqrt{x^2 + y^2}}) \end{cases}$$

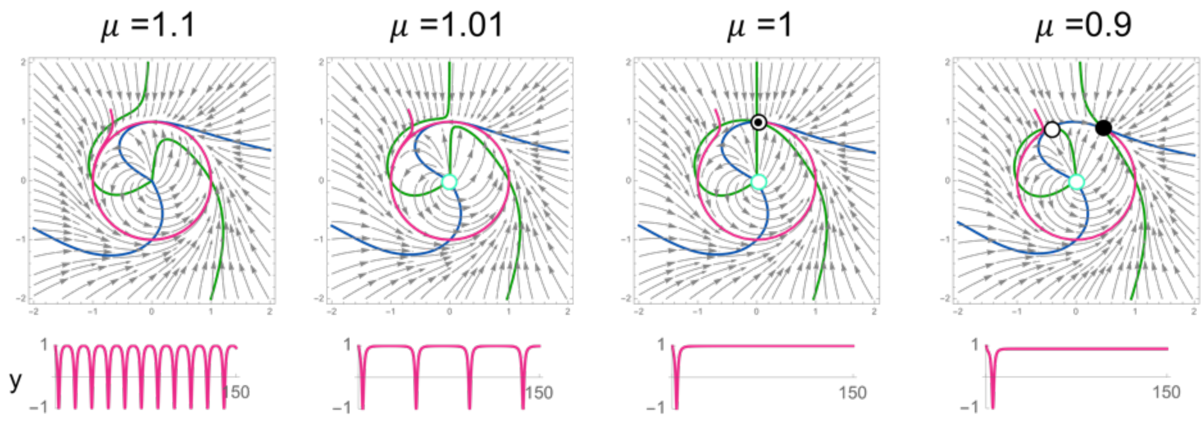

**Figure S13 SNIC bifurcation when varying $\mu$ around the bifurcation point**

### 5. Parameters optimization for 10 bits spike trains

This section describes how we optimized conditions to activate or deactivate on demand the oscillation of PP system with temperature. We optimized for several tradeoffs. First, the concentrations of species catalyzing the replication of prey (DNA template and polymerase) must not be set too low (because the concentrations of preys and their fluorescence would not be easily detectable), nor too high (because it makes it difficult to turn OFF the growth of preys)

(Figure S14). A polymerase activity of 50 U ml$^{-1}$ gives sustained oscillations but of low amplitude, while 75 U ml$^{-1}$ gives large damped oscillations. Therefore the midpoint of 62.5 U ml$^{-1}$ was chosen. As for template, a concentration of 60 nM of G1 template is a good compromise between high amplitude oscillations and the ability to turn the system off at high temperature.

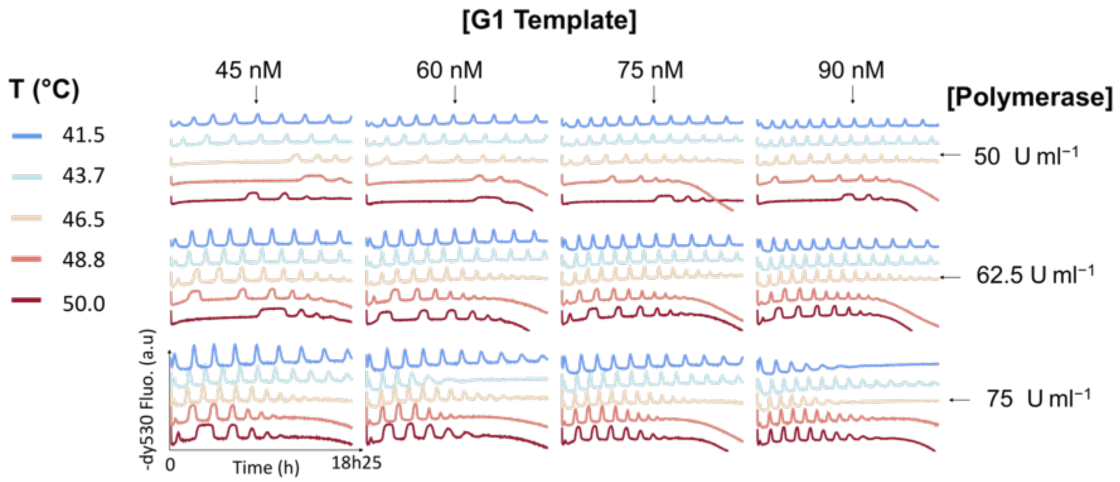

**Figure S14 - Optimization of the G1 template and polymerase with regard to temperature.** G1 ranging from 45 nM to 60 nM, N1 2 nM, P1 10 nM, Polymerase ranging from 50 U.mL$^{-1}$ to 75 U.mL$^{-1}$, Nickase 400 U.mL$^{-1}$, Exonuclease 27 nM, ETSSB 20 µg .mL$^{-1}$

Secondly, low concentrations of ETSSB slows the system down (because it takes more time to separate two predator strands during predation), whereas high concentrations of ETSSB promotes the emergence of parasitic species (because these parasitic strands need to separate their double stranded DNA [18], Fig. S15). Concentrations of ETSSB of 10 µg ml$^{-1}$ or lower give small and slow oscillations whereas parasitic species emerge sooner for concentrations higher than 15 µg ml$^{-1}$. ETSSB concentration was therefore fixed to 12.5 µg ml$^{-1}$ going forward.

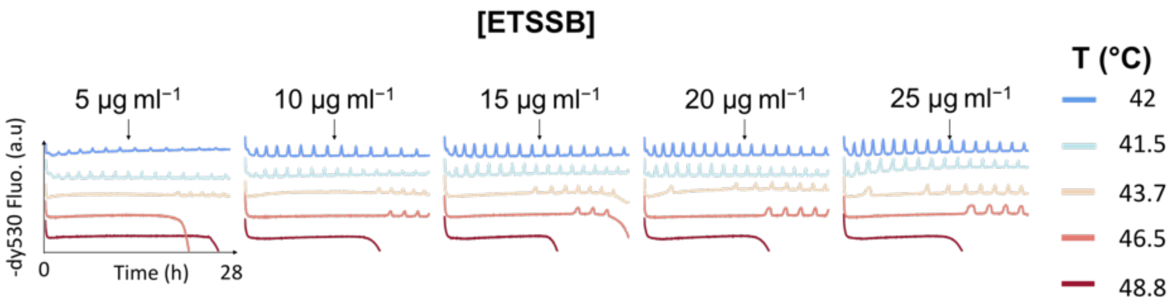

**Figure S15 - Optimization of the concentration of ETSSB.** G1 60 nM, N1 1 nM, P1 9 nM, Polymerase 62.5 U.mL$^{-1}$, Nickase 400 U.mL$^{-1}$, Exonuclease 30.6 nM, ESTSB ranging from 5 µg.mL$^{-1}$ to 25 µg.mL$^{-1}$

Lastly, the range of temperature must be chosen carefully (Figures S16 & S17). A large thermal amplitude between the OFF and ON temperatures made it easier to fall in the desired regimes. However this thermal amplitude can not be too large either. If the ON temperature is too low, the priming of the oscillations is slow (i.e. it takes more time to create the first batch of preys), making it hard to turn the system ON. At 41.8°C, the system does not oscillate fast enough to get enough peaks for messages with several consecutives “1”. The ON temperature was fixed at 43°C going forward.

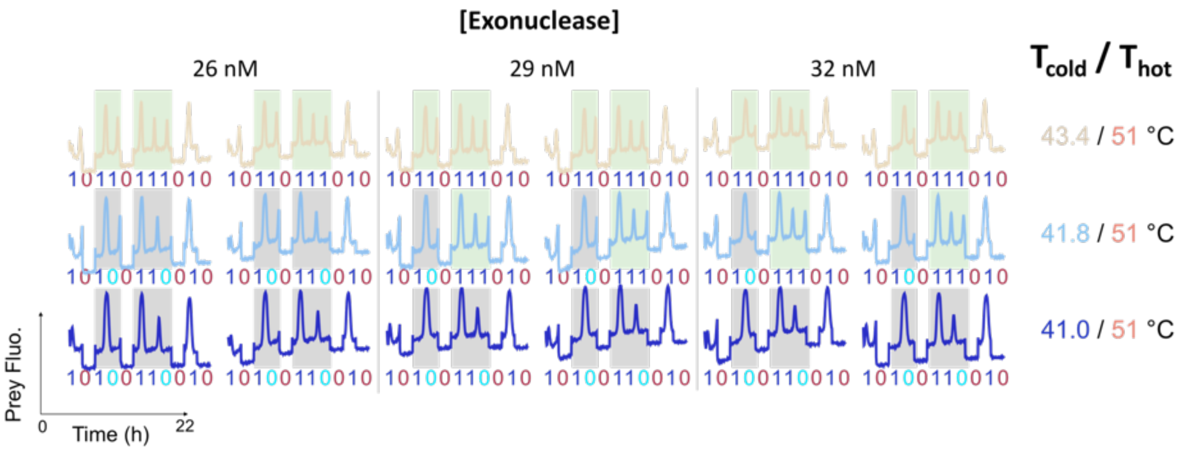

**Figure S16 - Optimization of the “cold temperature”.** Green rectangles denote successful transmission of sequences of 1 bits. G1 60 nM, N1 1 nM, P1 9 nM, Polymerase 62.5 U.mL$^{-1}$, Nickase 400 U.mL$^{-1}$, Exonuclease 29 nM, ESTSB 12.5 µg mL$^{-1}$ (same composition as Figure 1D)

The OFF temperature needs to be chose above the bifurcation ~51°C, but high temperatures promote the emergence of parasitic species (likely due to temperature-assisted partial melting of parasitic strands[18]). Temperatures lower than 52°C could not transmit long series of “0” bits without spiking. The OFF was fixed at 52°C going forward.

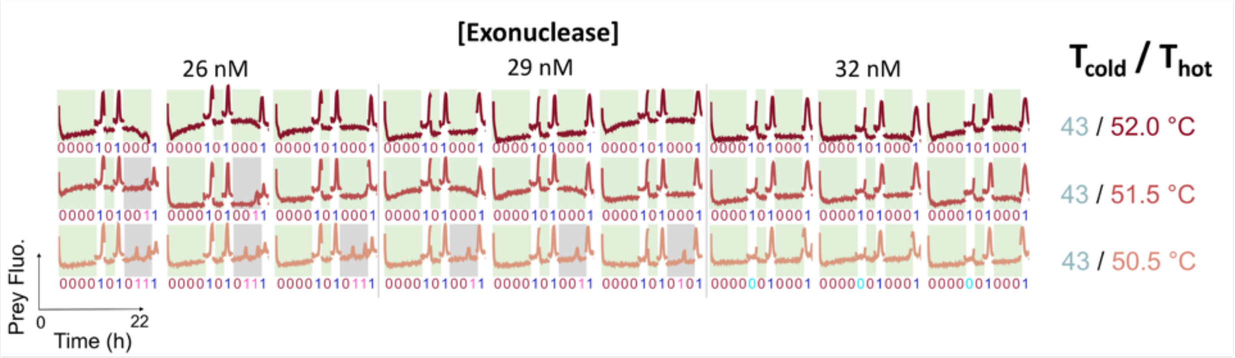

**Figure S17 - Optimization of the “hot temperature”.** Green rectangles denote successful transmission of sequences of 0 bits. G1 60 nM, N1 1 nM, P1 9 nM, Polymerase 62.5 U.mL$^{-1}$, Nickase 400 U.mL$^{-1}$, Exonuclease 29 nM, ESTSB 12.5 µg mL$^{-1}$ (same composition as Fig 1D)
