## Supplementary material for "Neural coding of temperature with a DNA-based spiking chemical neuron": READ ME - Videos & Image Legends

### Movie S1 Legend:

2D phase portraits of the chemical neuron for decreasing temperatures (See Fig 2A). Thick green and blue lines are the nullclines for preys and predators respectively (see 2.1). The thin pink line shows a generic trajectory. Steady states – located at the intersection of nullclines – are shown as disks, color-coded by their nature and stability (see SI 2.6)

### Movie S2 Legend:

2D phase portraits of a toy model of biological thermosensory neuron (see Fig 2B), which has 2 variables (membrane potential x and spiking variable y), and is parametrized by temperature through its excitation current (see SI 3.1). The blue curve is the nullcline of the membrane potential, and the green curve is the nullcline of the spiking variable. The thin pink line shows a generic trajectory. Steady states – located at the intersection of nullclines – are shown as disks, color-coded by their nature and stability (see SI 2.6)

### Movie S3 Legend:

Time-lapse of encapsulated chemical neurons in a temperature gradient (see Fig. 6) for different exonuclease concentrations given by a fluorescent barcoding (Image S1).

#### Image S1 Legend:

Fluorescence barcoding allowing to assess the concentration of exonuclease in each droplet. Concentration of exonuclease increases from dark blue (15nM) to white (45 nM)
