## Supplementary figures and images for "Neural coding of temperature with a DNA-based spiking chemical neuron"

### Image S1-Barcoding of Exonuclease concentration in Encapsulated Chemical neurons

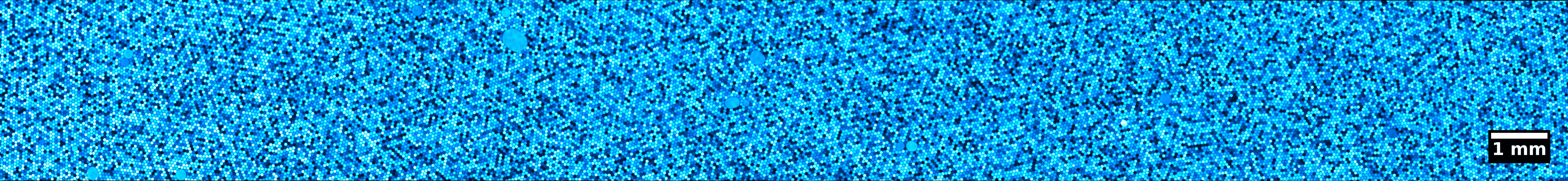
